# DeepSleep: Fast and Accurate Delineation of Sleep Arousals at Millisecond Resolution by Deep Learning

**DOI:** 10.1101/859256

**Authors:** Hongyang Li, Yuanfang Guan

## Abstract

Sleep arousals are transient periods of wakefulness punctuated into sleep. Excessive sleep arousals are associated with many negative effects including daytime sleepiness and sleep disorders. High-quality annotation of polysomnographic recordings is crucial for the diagnosis of sleep arousal disorders. Currently, sleep arousals are mainly annotated by human experts through looking at millions of data points manually, which requires considerable time and effort. Here we present a deep learning approach, DeepSleep, which ranked first in the 2018 PhysioNet Challenge for automatically segmenting sleep arousal regions based on polysomnographic recordings. DeepSleep features accurate (area under receiver operating characteristic curve of 0.93), high-resolution (5-millisecond resolution), and fast (10 seconds per sleep record) delineation of sleep arousals.

## Main

Sleep is important for our overall health and quality of life ^1^. Inadequate sleep is often associated with many negative outcomes, including obesity ^2^, irritability ^2,3^, cardiovascular dysfunction ^4^, hypotension ^5^, impaired memory ^6^ and depression ^7^. About one third of the general population in the United States are affected by insufficient sleep ^8^. The prevalence of inadequate sleep results in large economic costs ^9^ and continues to increase in various nations ^10,11^. Spontaneous sleep arousals, defined as brief intrusions of wakefulness into sleep ^12^, are a common characteristic of brain activity during sleep. Excessive arousals due to disturbances can be harmful, resulting in fragmented sleep, daytime sleepiness and sleep disorders ^13,14^. There are different types of arousing ^15^stimulus, including obstructive sleep apneas or hypopneas, respiratory effort-related arousals (RERA), hyperventilations, bruxisms (teeth grinding), snoring, vocalizations, and leg movements. Together with sleep stages (wakefulness, stage1, stage2, stage3, and rapid eye movement), sleep arousals are labeled through visual inspections of polysomnographic recordings according to the American Academy of Sleep Medicine (AASM) scoring manual ^16^. Of note, an 8-hour sleep record sampled at 200Hz with 13 different physiological measurements contains a total of 75 million data points. It takes hours to manually annotate such a large-scale sleep record.

Many research efforts have been made in developing computational methods for automatic arousal detection based on polysomnographic recordings ^17–21^. These methods mainly focus on 30-second epochs, and extract statistical features in the time and frequency domains through Fourier transform or in-house feature engineering. These features and/or raw signals are subsequently fed into machine learning models to predict sleep arousals. However, due to the large differences of datasets and evaluation metrics used in previous studies, it remains unknown how to build an accurate and robust model to quickly delineate all sleep arousal events within a sleep record at a high resolution. In particular, how to preprocess the raw data or extract features before training models? Which types of machine learning models are well suited? What is the optimal input length (e.g. 30-second epochs or full-length records)? Which types of physiological signals should be used?

Here we investigate these questions and describe a novel deep learning approach, DeepSleep, for automatic detection of sleep arousals. This approach ranked first in the 2018 “You Snooze, You Win” PhysioNet/Computing in Cardiology Challenge ^22^, in which state-of-the-art computational methods were systematically evaluated for predicting non-apnea sleep arousals on a large held-out test dataset ^23^. The workflow of DeepSleep is schematically illustrated in **Fig. 1**. We built a deep convolutional neural network (CNN) to capture long-range and short-range interdependencies between time points across an entire sleep record. Information at different resolutions and scales was integrated to improve the performance. Intriguingly, we found that similar EEG and EMG channels were interchangeable, which was used as a special augmentation in our approach. DeepSleep is able to delineate the sleep arousal profile of a sleep record at 5-millisecond resolution within 10 seconds.

**Fig. 1.**
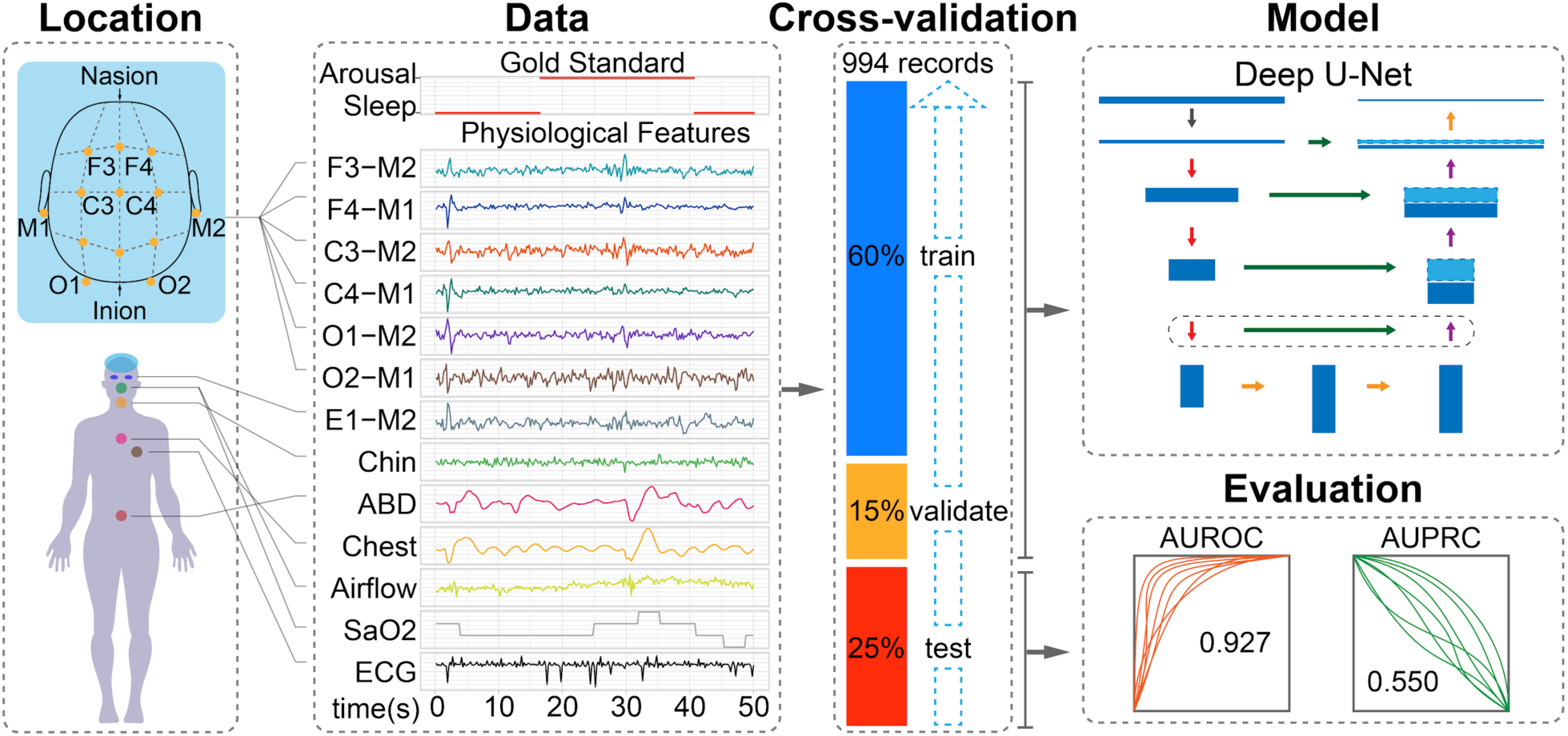
Schematic Illustration of DeepSleep workflow. **Location.** The 13-channel polysomnogram monitored multiple body functions, including brain activity (EEG), eye movement (EOG), muscle activity (EMG), and heartbeat (ECG). **Data.** A 50-second sleep record with the gold standard label of arousal/sleep and 13 physiological features. **Cross-validation.** In the nested train-validate-test framework, 60%, 15%, and 25% of the data were used to train, validate, and evaluate the model. **Model.** The classic U-Net architecture was adapted to capture the information at different scales and allowed for detecting sleep arousals at millisecond resolution. **Evaluation.** DeepSleep achieved high area under receiver operating characteristic curve (AUROC) of 0.927 and area under precision-recall curve (AUPRC) of 0.550 on the testing dataset.

## Results

### Overview of the experimental design for predicting sleep arousals from polysomnogram

In this work, we used the 994 polysomnographic records provided in the 2018 PhysioNet challenge, which were collected at the Massachusetts General Hospital. In each record, 13 physiological measurements were sampled at 200Hz (Location and Data in **Fig. 1**), including six electroencephalography (EEG) signals at F3-M2, F4-M1, C3-M2, C4-M1, O1-M2 and O2-M1; one electrooculography (EOG) signal at E1-M2; three electromyography (EMG) signals of chin, abdominal and chest movements; one measure of respiratory airflow; one measure of oxygen saturation (SaO_2_); one electrocardiogram (ECG). Each time point in the polysomnographic record was labeled as “Arousal” or “Sleep” by sleep experts, excluding some non-scoring regions such as apnea or hypopnea arousals. To exploit the information of the training records, we employed a nested train-validate-test framework, in which 60% of the data was used to train the neural network, 15% of the data was used to validate for parameter selection and 25% of the data was used to evaluate the performance of the model (Cross-validation in **Fig. 1**). To capture the long-range and short-range information at different scales, we adapted a classic neural network (Model in **Fig. 1**), U-Net, which was originally designed for image segmentation ^24^. Multiple data augmentation strategies, including swapping similar polysomnographic channels, were used to expand the training data space and enable the generalizability of the model. Finally, the prediction performance was evaluated by the area under receiver operating characteristic curve (AUROC) and the area under precision-recall curve (AUPRC) on the held-out test dataset of 989 records (Evaluation in **Fig. 1**) during the challenge.

### Highly heterogeneous sleep records among individuals

By investigating the annotations of these sleep records, we found high levels of heterogeneity among individuals. In **Fig. 2A**, we randomly selected sleep records of 20 individuals and presented the annotations in different colors. There are 8 major annotation categories: “Arousal”, “Undefined”, “REM” (Rapid Eye Movement), “N1” (Non-REM stage 1), “N2” (Non-REM stage 2), “N3” (Non-REM stage 3), “Wake” and “Apnea”. The distribution of these categories differs dramatically among individuals (different colors in **Fig. 2A**). Clearly, different individuals display distinct patterns of sleep, including the length of total sleep time and multiple sleep stages. Notably, the sleep arousal regions are relatively short and sparsely distributed along the entire record for most individuals (yellow regions in **Fig. 2A**).

**Fig. 2.**
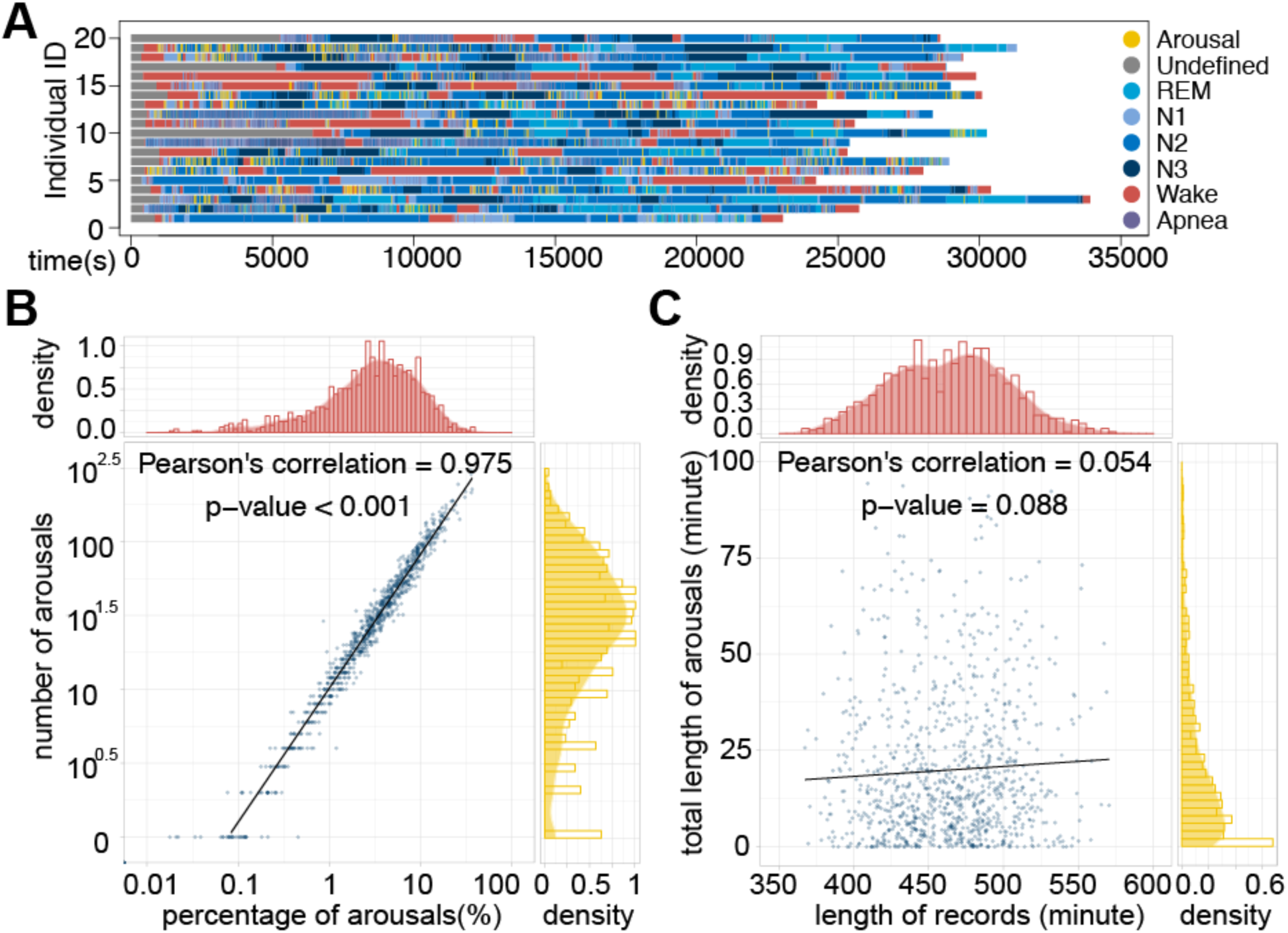
Sleep arousals sparsely distributed in the heterogenous sleep records among individuals. (**A**) The 8 major annotation categories are shown in different colors for 20 randomly selected sleep records. The apneic and non-apneic arousal events overwrite sleep stages (N1, N2, N3, REM). (**B**) The relationship between the number of sleep arousals (y-axis) and the percentage of total sleep arousal time over total sleep time (x-axis) in the 994 sleep records. In general, more arousal events lead to longer accumulated arousal time and the correlation is significantly strong. (**C**) The length of sleep (x-axis) has no significant correlation with the accumulated length of sleep arousals (y-axis).

We further investigated the occurrence of arousals and found that the median number of arousals during sleep was 29, indicating the prevalence of sleep arousals. A total of 43 individuals (4.33%) had solid sleep without any arousal, whereas 82 individuals (8.25%) had more than 100 arousals during their sleep (y-axis in **Fig. 2B)**, lasting around 10% of the total sleep duration (x-axis in **Fig. 2B**). In addition, there was no significant correlation between the total sleep time and the total length of sleep arousals (**Fig. 2C**), which was expected since the quality of sleep is not determined by sleep length. In summary, the intrinsically high heterogeneity of sleep records across individuals rendered the segmentation of sleep arousals a very difficult problem.

### Deep U-Net captures the long-range and short-range information at different scales and resolutions

Current manual annotation of sleep arousals is defined by the AASM scoring manual ^16^, in which sleep experts focus on a short period (less than a minute) and make decisions about sleep arousal events. However, it remains unclear whether the determinants of sleep arousals reside only within a short range, or long-range information across minutes and even hours plays an indispensable role in detecting sleep arousals. Although sleep arousal is in nature a transient event, it may be associated with the overall sleep pattern through the night. Intriguingly, when we trained the convolutional neural networks on longer sleep records, we consistently achieved better performances (**Fig. S1**). Therefore, we used the entire sleep record as input to make predictions, instead of small segments of a sleep record.

To learn the long-range association between data points across different time scales (second, minute, and hours), we develop an extremely deep convolutional neural network, which contains a total of 35 convolutional layers (**Fig. 3A**). This network architecture has two major components, the encoder and the decoder. The encoder takes a full-length sleep record of 2^23^ = 8,388,608 time points and gradually encrypts the information into a latent space (the red trapezoid in **Fig. 3A**). Sleep recordings were centered, regardless of their original lengths, within the 8-million input space by filling in with zeros on their extremes. To be specific, the convolution-convolution-pooling (hereafter referred to as “ccp”) block is used to gradually reduce the size from 2^23^ = 8,388,608 to 2^8^ = 256 (**Fig. 3B** top). Meanwhile, the number of channels gradually increases from 13 to 480 to encode more information, compensating the loss of resolution in the time domain. In each convolutional layer, the convolution operation is applied on the data along the time axis to aggregate the neighborhood information. Since the sizes of data in these convolutional layers are different, the encoded information is unique within each layer. For example, in the input layer, 10 successive time points sampled at 200Hz correspond to a short time interval of 10/200=0.05 seconds, whereas in the center layer (size = 2^8^), 10 time points correspond to a much longer time interval of 0.05 * 2^23-8^ = 1,638 seconds, nearly 30 minutes. Therefore, this deep encoder architecture allows us to capture and learn about the interactions across data points at multiple time scales. The relationship between length of segments and the corresponding time can be found in **Table S1**.

**Fig. 3.**
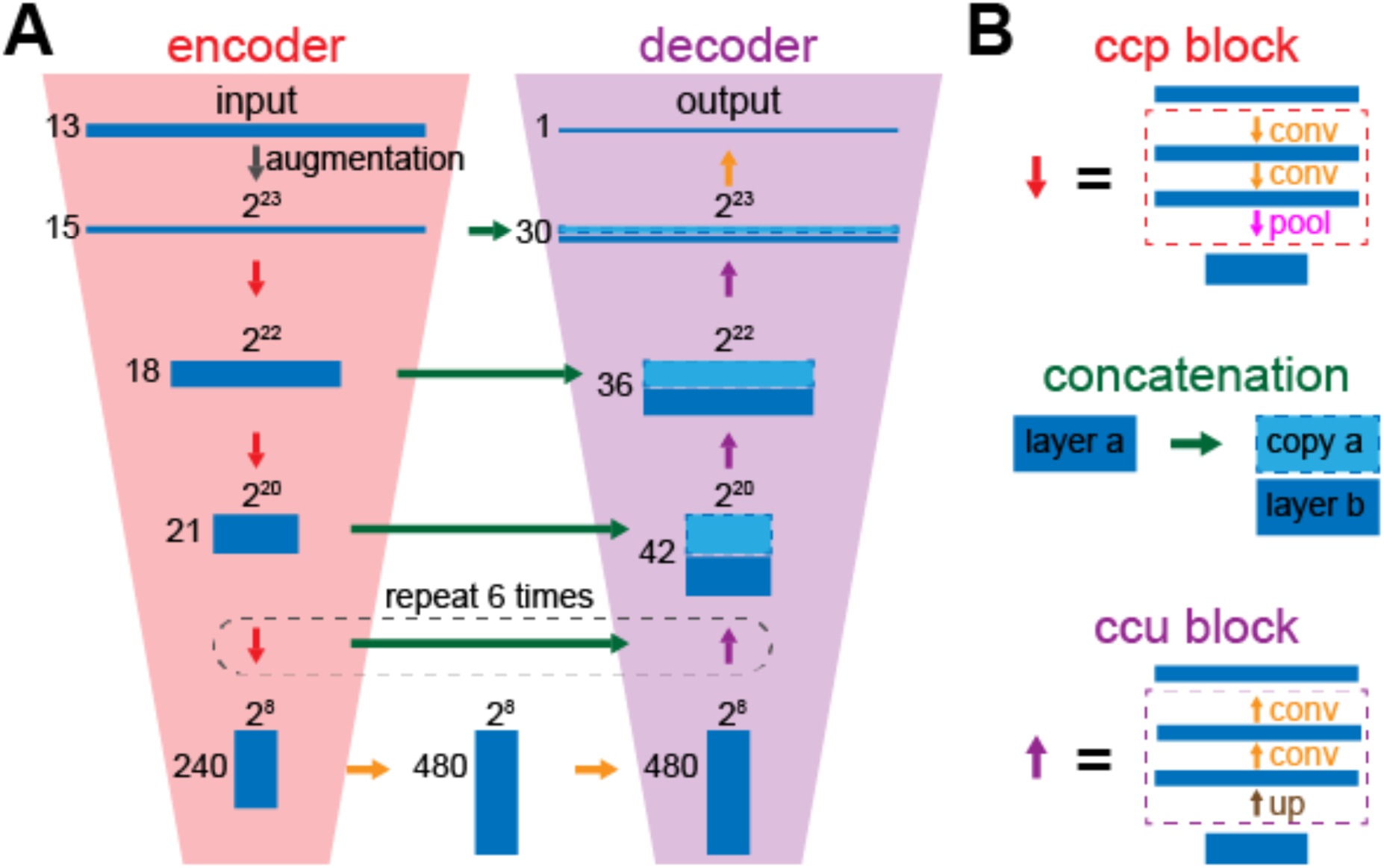
The deep convolutional neural network architecture in DeepSleep. (**A**) The classic U-Net structure was adapted in DeepSleep, which has two major components of the encoder (the red trapezoid on the left) and the decoder (the purple trapezoid on the right). (**B**) The building blocks of DeepSleep are the convolution-convolution-pooling block (red), the concatenation (green) and the convolution-convolution-upscaling block (purple). The orange arrow represents the convolution operation.

Similar to the encoder, the second component of our network architecture is a decoder to decrypt the compressed information from the center latent space. In contrast to the “ccp” block, the convolution-convolution-upscaling (hereafter referred to as “ccu”) block is used (**Fig. 3B** bottom), which gradually increases the size and decreases the number of channels of the data (the purple trapezoid in **Fig. 3A**). In addition, concatenation is used to integrate the information from both the encoder and the decoder at each time scale (green horizontal arrows in **Fig. 3**). Finally, the output is the segmentation of the entire sleep record, where high prediction values indicate sleep arousal events and low values indicate sleep.

### Deep learning enables accurate predictions of sleep arousals

By capturing the information at multiple resolutions, DeepSleep achieves high performance in automatic segmentation of sleep arousals. Since deep neural networks are iteration-based machine learning approaches, a validation subset is used for monitoring the underfitting or overfitting status of a model and approximating the generalization ability on unseen datasets. A subset of 15% randomly selected records was used as the validation set during the training process (Cross-validation in **Fig. 1**) and the cross entropy was used to measure the training and validation losses (see details in **Materials and Methods**). The 13 polysomnographic channels complemented each other and using all of them instead of one type of these signals enabled the neural network to capture interactions between channels and achieved the highest performance (**Fig. S2A-B**). We developed three basic models called “1/8”, “1/2” and “full”, according to the resolution of the neural network input. The “full” resolution means that the original 8-million (2^23^ = 8,388,608) length data were used as input. The “1/2” or “1/8” resolution means that the original input data were first shrunk to the length of 4-million (2^22^) or 1-million (2^20^) by averaging every 2 or 8 successive time points, respectively. We observed similar validation losses of the “full”, “1/2” and “1/8” models (solid lines in **Fig. 4A**). The final evaluation was based on the AUROC and AUPRC scores of predicting 25% of the data. In **Fig. 4B**, each blue dot represented one sleep record and we observed a significant yet weak correlation = 0.308 between the AUROCs and AUPRCs. The baselines of random predictions were shown as red dashed lines. Notably, the AUPRC baseline of 0.072 corresponded to the ratio of the average total sleep arousal length over the total sleep time, which was considerably low and made it a hard task due to the intrinsic sparsity of sleep arousal events.

**Fig. 4.**
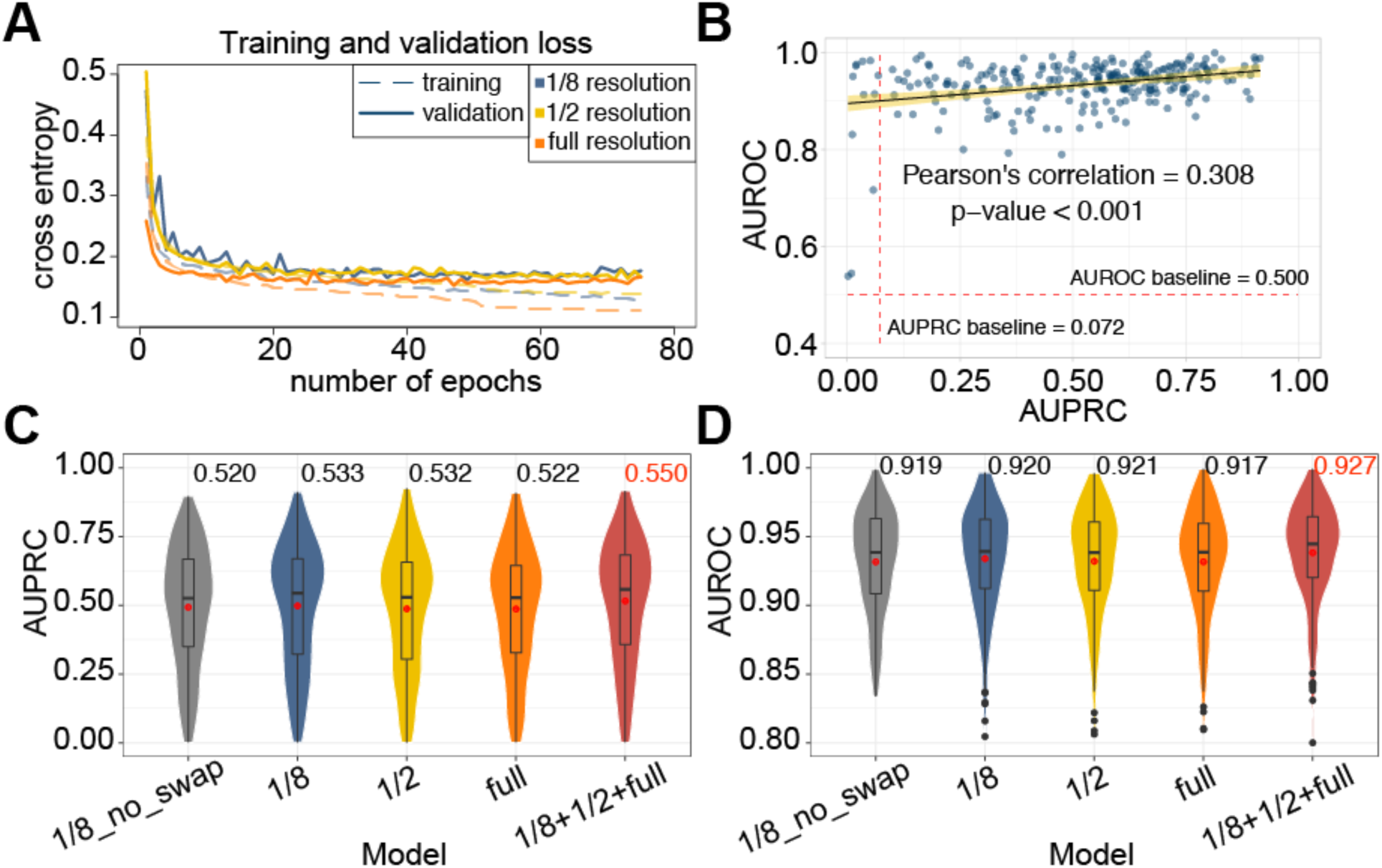
The performance comparison of DeepSleep using different model training strategies. (**A**) The training and validation cross entropy losses are shown in the dashed and solid lines, respectively. The models using sleep records at different resolutions are shown in different colors. (**B**) The prediction of each sleep record in the test set is shown as a blue dot in the AUROC-AUPRC space. A weak correlation is observed between AUROCs and AUPRCs with a significant p-value < 0.001. The 95% percent confidence interval is shown as the yellow bend. The baselines of random predictions are shown as red dashed lines. The prediction (**C**) AUPRCs and (**D**) AUROCs of models using different resolution or strategies were calculated. The “1/8_no_swap” model corresponds to the model using the “1/8” resolution records as input without any channel swapping, whereas the “1/8”, “1/2” and “full” models use the strategy of swapping similar polysomnographic channels. The final “1/8+1/2+full” model of DeepSleep is the ensemble of models at 3 different resolutions, achieving the highest AUPRC of 0.550 and AUROC of 0.927.

To build a robust and generalizable model, multiple data augmentation strategies were used in DeepSleep. After carefully examining the data, we found that signals belonging to the same physiological categories were very similar and synchronized, including two EMG channels and six EEG channels (see Data in **Fig. 1**). We applied a novel augmentation strategy by randomly swapping these similar channels during the model training process, assuming that these signals were interchangeable in determining sleeping arousals. There are three EMG channels but EMG-chin were not considered in this swapping strategy due to its differences from the other two EMG (ABD and chest) channels (see Data in **Fig. 1**). This channel swapping strategy was bold but effective, adapting which largely improved the prediction performance (“1/8_no_swap” versus “1/8” in **Fig. 4C-D**). In addition, we multiplied the polysomnographic signals by a random number between 0.90 and 1.15 to simulate the inherent fluctuation and noise of the data. Other augmentations on the magnitude and time scale were also explored (**Fig. S2C-D**). Furthermore, to address the heterogeneity and batch effects among individuals, we quantile normalized each sleep record to a reference, which was generated by averaging all the records. This step effectively removed the biases introduced by the differences of individuals and instruments, and Gaussian normalization was also tested and had slightly lower performance (**Fig. S2E-F**). Finally, we assembled the predictions from the “1/8”, “1/2” and “full” resolution models as the final prediction in DeepSleep (red violin plots in **Fig. 4C-D**).

We further compared different machine learning models and strategies in segmenting sleep arousals. We first tested a classical model, logistic regression, and found that our deep learning approach had a much higher performance (**Fig. S2G-H**). It has also been reported that neural network approaches significantly outperformed classical machine learning methods, including random forest, logistic regression ^25^, support vector machine, and linear models ^26^. In fact, 8 out of the top 10 teams used neural network models in the 2018 PhysioNet Challenge (red blocks in **Fig. S3A**) ^22^. Two types of network structures (convolutional and recurrent) were mainly used, and integrating Long Short-Term Memory (LSTM) or Gated Recurrent Unit (GRU) into DeepSleep did not improve the performance (**Fig. S3B-D**). In terms of input length, increasing input length significantly improved the performance, and full-length records were used by three teams (blue blocks in **Fig. S3A**). We also compared DeepSleep with recent state-of-the-art methods in sleep stage scoring. These methods extracted features from 30-second epochs through short-time Fourier transform (STFT) ^27,28^ or Thomson’s multitaper ^25,29^. They were originally designed for automatic sleep staging and we applied them to the task of detecting sleep arousals on the same 2018 PhysioNet data. Although these methods performed well in sleep stage scoring, they were not well suited for arousal detection (**Fig. S3E-F**). Deep learning approaches can model informative features in an implicit way without tedious feature crafting ^30^, and neural networks using raw data as input were frequently used by half of the top 10 teams (orange blocks in **Fig. S3A**).

To comprehensively investigate the effects of various network structures and parameters on predictions, we further performed experiments with different modifications, including shallow neural network (**Fig. S4A-B**), average pooling (**Fig. S4C-D**), large convolution kernel size (**Fig. S4E-F**), and loss functions (**Fig. S4G-H**). These modifications had either similar or lower prediction performances. We concluded that the neural network architecture and augmentation strategies in DeepSleep were optimized for the current task of segmenting sleep arousals. Subsequent analysis of the relationships between the prediction performance and the number of arousal were investigated (**Fig. S5A-B**). As we expected, the prediction AUPRC was correlated with the number of arousals in a sleep record. The individuals who had more sleep arousals during sleep were relatively easier to predict. Moreover, we tested the runtime of DeepSleep with Graphics Processing Unit (GPU) acceleration and segmenting sleep arousals of a full sleep record can be finished within 10 seconds on average (**Fig. S5C-D**). The time cost of DeepSleep is much lower than that of manual annotations, which requires hours for one sleep record.

In addition to the 2018 PhysioNet dataset, we further validated our method on the large publicly available Sleep Heart Health Study (SHHS) dataset, which contains 6,441 individuals in SHHS visit 1 (SHHS1) and 3,295 individuals in SHHS visit 2 (SHHS2) ^31^. The SHHS is a multi-center cohort study, including participants from multiple different cohorts and the polysomnograms were annotated by sleep experts from different labs (https://sleepdata.org/datasets/shhs). The recording montages and signal sampling rates of SHHS1 and SHHS2 were quite different. For both SHHS1 and SHHS2, we randomly selected 1,000 recordings, which was comparable to the number of recordings (n=994) in the PhysioNet training dataset. Then we applied DeepSleep pipeline to train, validate, and test models on SHHS1 and SHHS2 datasets individually. We observed similar performances of detecting sleep arousals on the PhysioNet, SHHS1, and SHHS2 datasets in **Fig. S6A-B**, demonstrating the robustness of our DeepSleep method.

In the clinical setting, both apneic and non-apneic arousal are very important. We have therefore built neural network models for detecting apnea, in addition to the model for detecting non-apneic arousals, which was originally designed during the 2018 PhysioNet challenge. Specifically, we applied DeepSleep pipeline to the PhysioNet data and built three types of models for (1) detecting apneic arousals; (2) detecting non-apneic arousals; and (3) detecting all arousals (apneic and non-apneic arousals). DeepSleep is able to detect both apneic and non-apneic arousals (**Fig. S6C-D**).

### Visualization of DeepSleep predictions

In addition to the abstract AUROC and AUPRC scores, we directly visualized the prediction performance of DeepSeep at 5-millisecond resolution (corresponding to the 200Hz sample rate). An example 7.5-hour sleep record with the prediction AUROC of 0.960 and AUPRC of 0.761 is shown in **Fig. 5**. More examples at 3 rank percentiles (25%, 50%, and 75%) based on the AUPRC values can be found in **Fig. S7**. From top to bottom, we plotted the multi-stage annotations, sleep arousal labels, predictions and cross-entropy losses long the time x-axis. By comparing the prediction and gold standard, we can see the general prediction pattern of DeepSleep correlates well with the gold standard across the entire record (the second and third rows in **Fig. 5A**). We further zoom into a short interval of 12.5 minutes and DeepSleep successfully identifies and segments seven sleep arousal events out of eight (yellow in **Fig. 5B**), although one arousal around 25,600 is missed. Intriguingly, DeepSleep predictions display a different pattern from the gold standard annotated by sleep experts: DeepSleep assigns continuous prediction values with lower probabilities near the arousal-sleep boundaries, whereas the gold standard is strictly binary either arousal = 1 or sleep = 0 based on the AASM scoring manual ^16^. This becomes clearer when examining the cross entropy loss at each time point and the boundary region has higher losses shown in red (the bottom row in **Fig. 5B**). This is expected because in general we will have a higher confidence of annotation in the central region of sleep arousal or other sleep events. Yet due to the limit of time and effort, it is practically infeasible to introduce rules for manually annotating each time point via a probability scenario. Additionally, binary annotation of sleep records containing millions of data points has already required significant effort. DeepSleep opens a new avenue to reconsider the way of defining sleep arousals or other sleep stage annotations by introducing the probability system.

**Fig. 5.**
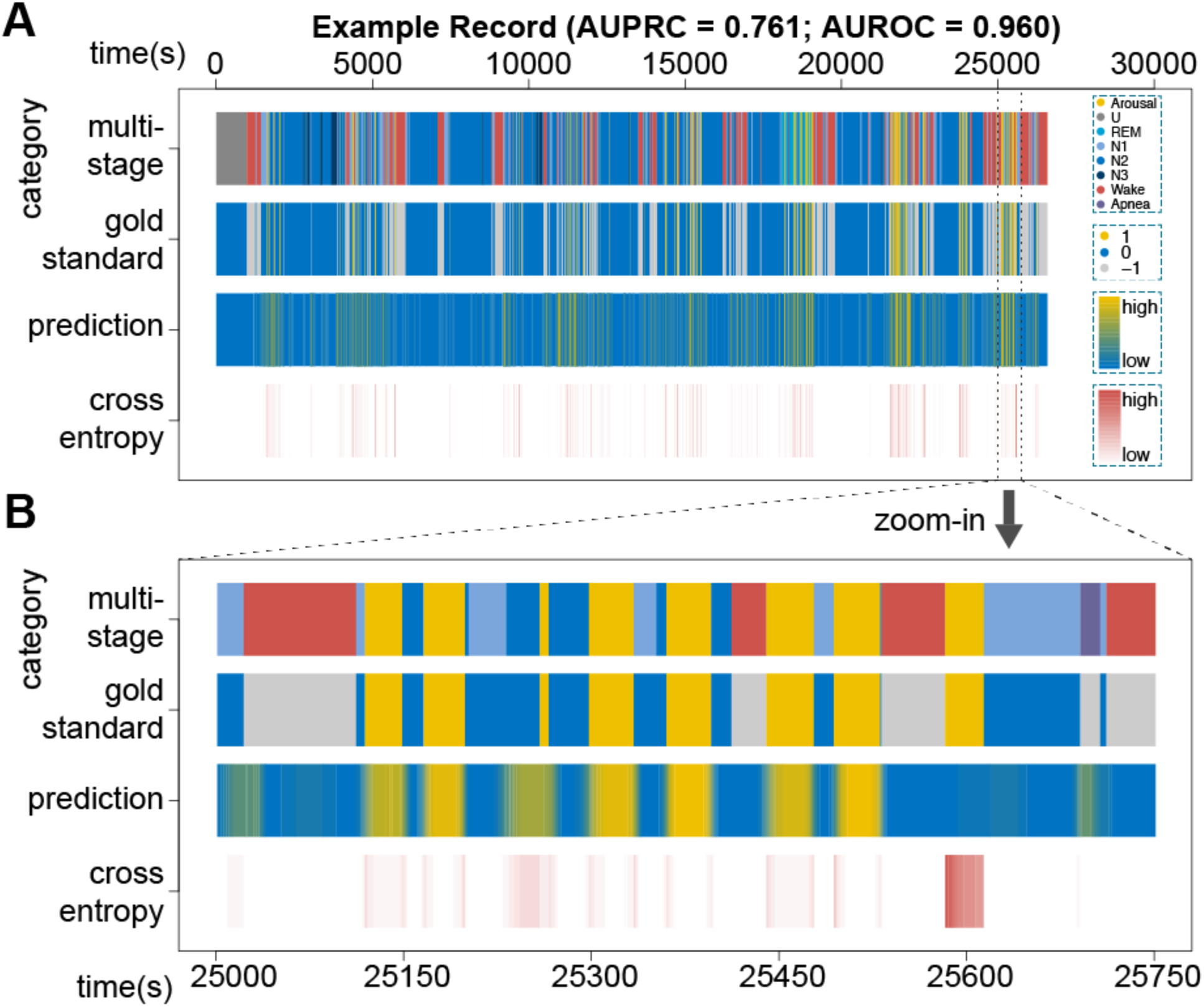
Visualization of DeepSleep predictions and the gold standard annotations. (**A**) A 7.5-hour sleep record (id=tr05-1034) with the prediction AUROC of 0.960 and AUPRC of 0.761 is used as an example. From top to bottom along the y-axis, the four rows correspond to the 8 annotation categories, the binary label of arousal (yellow), sleep (blue) and the non-scoring regions (gray), the continuous prediction, and the cross-entropy loss at each time point along the x-axis. The wrongly predicted regions lead to high cross entropy losses, which are shown in dark red at the bottom row. (**B**) The zoomed in comparison of a 12.5-minute period of this sleep record.

## Discussion

In this study, we created a deep learning approach, DeepSleep, to automatically segment sleep arousal regions in a sleep record based on the corresponding polysomnographic signals. A deep convolutional neural network architecture was designed to capture the long-range and short-range interactions between data points at different time scales and resolutions. Unlike classical machine learning models ^32^, deep learning approaches do not depend on manually crafted features and can automatically extract information from large datasets in an implicit way ^33^. Using classical approaches to define rules and craft features for modelling sleep problems in real life would become much too tedious. In contrast, without assumptions and restrictions, deep neural networks can approximate complex mathematical functions and models to address those problems. Currently, these powerful tools have also been successfully applied to biomedical image analysis and signal processing ^34,35^. Compared with classical machine learning models, deep learning is a “black box” method which is relatively hard to interpret and understand. Meanwhile, deep learning approaches usually requires more computational resources such as GPUs, whereas most classical machine learning models can run on common CPUs.

Overfitting is a common issue in deep learning models, especially when the training dataset is small and the model is complex. Even if we use a large dataset and perform cross-validation, we will gradually and eventually overfit to the data. This is because each time we evaluate a model using the internal test set, we probe the dataset and fit our model to it. In contrast to previous studies, the 2018 PhysioNet Challenge offered us a unique opportunity to truly evaluate the performances and compare cutting-edge methods on a large external hidden test set of 989 samples ^23^. In addition, we demonstrate that deep convolutional neural networks trained on full-length records and multiple physiological channels have the best performance in detecting sleep arousals, which are quite different from current approaches extracting features from short 30-second epochs ^25,27,30^. Beyond sleep arousals, we propose that the U-Net architecture used in DeepSleep can be adapted to other segmentation tasks such as sleep staging. A multi-tasking learning approach can be further implemented as the outputs of U-Net to directly segment multiple sleep stages simultaneously based on polysomnograms.

An interesting observation is that when we used records of different lengths as input to train deep learning models, the model using full-length records largely outperformed models using short periods of records. This observation brings about the question of how to accurately detect sleep arousals based on polysomnography. Current standards mainly focus on short time intervals of less than one minute ^16^, yet the segmentations among different sleep experts are not very consistent in determining sleep arousals. One reason is that it is hard for humans to directly read and process millions of data points at once. In contrast, computers are good at processing large-scale data and discover the intricate interactions and structures between data points across seconds, minutes and even hours. Our results indicate that sleep arousal events are not be solely determined by the local physiological signals but associated with much longer time intervals even spanning hours. It would be interesting to foresee the integration of computer-assisted annotations to improve definitions of sleep arousals or other sleep stages.

In addition to the unique long-range information captured by DeepSleep, a clear advantage of computational approaches lies in the annotations for the boundary regions between arousal and sleep. Since current sleep annotations are binary only, it would be a more accurate and appropriate approach to introduce the probability of the annotation confidence, especially at the boundary regions. Machine learning approaches such as DeepSleep naturally provide the continuous predictions for each time point. It would be interesting to see improved annotation systems using continuous values instead of binary labels. A simple approach could be directly integrating the computer predictions with annotations by human sleep experts. The proposed annotation systems would provide more accurate information for the diagnosis of sleep disorders and the evaluation of sleep quality in the future.

## Materials and Methods

### Polysomnographic recordings

The dataset used in this study contains a total of 994 polysomnographic sleep records from different individuals and their corresponding labels at each time point. Specifically, the arousal region is labeled by “1” and other sleep regions are labeled by “0”, except for the wakefulness regions, apnea arousal regions and hypopnea arousal regions labeled by “-1”. These “-1” regions will not be scored in the challenge, and we mainly focused on non-apnea arousals that interrupted the sleep of an individual, including spontaneous arousals, respiratory effort related arousals (RERA), bruxisms, hypoventilations, hypopneas, apneas (central, obstructive and mixed), vocalizations, snores, periodic leg movements, Cheyne-Stokes breathing or partial airway obstructions (https://physionet.org/challenge/2018/). The final test dataset consists of 989 unseen polysomnographic recordings from different individuals. For each time point sampled at 200Hz in each test sleep record, the participants needed to provide a prediction value between 0 and 1. A 8-hour sleep record contained nearly 75 million data points (8*60*60*200*13=74,880,000). Our model made predictions for all the time points, at the resolution of 5 milliseconds (1/200Hz = 5 milliseconds).

### Partition of the training, validation and testing sleep records

The 994 sleep records were randomly partitioned into three sets: 60% of them as the training set, 15% of them as the validation set and 25% of them as the testing set. The validation set was used for monitoring the training-validation losses and avoiding the problems of overfitting or underfitting.

### Gaussian normalization

The Gaussian normalization is calculated by

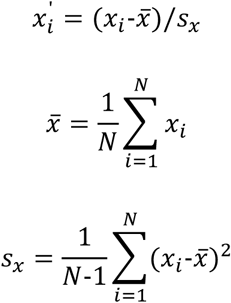

where x_i_ is the original value at time point i, x^’^_i_ is the normalized value at time point i, and N is the total number of time points. For the polysomnographic signals, we normalized each channel individually.

### Quantile normalization

For each polysomnographic channel, we first ranked the original input vector

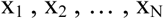

into a sorted vector in the increasing order

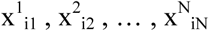

where superscript number denotes the ranked increasing order, and the subscript number denotes the original position before ranking. Then we replace this sorted vector with a sorted reference vector

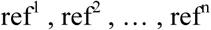

which is also in the increasing order. For example, x^k^_ik_ will be replaced by ref^k^. Then we changed the order back and mapped ref^k^ to its original position ik. After this quantile normalization, the overall distribution of the input vector has been mapped to the distribution of the reference vector. The reference vector was pre-calculated by averaging all the sorted recordings in the training dataset. We quantile normalized each recording to the same reference to address potential batch and cohort effects. Each polysomnographic channel was normalized individually.

### AUROC and AUPRC

Since sleep arousal events are extremely rare (<10% in terms of length), the performances of different methods are not apparent in the Receiver Operating Characteristic (ROC) curve, where the y-axis is the True Positive Rate (TPR) and the x-axis is the False Positive Rate (FPR). The TPR and FPR are defined as

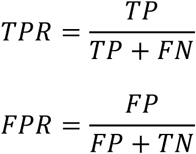

where TP is True Positive, FN is False Negative, FP is False Positive, and TN is True Negative. This is because when the number of negative events (“Sleep”; 92.8%), or TN, is much larger than the positive ones (“Arousal”; 7.2%), the FPR is always very small and will barely change even if a poor model makes many FP predictions. Therefore, in addition to the commonly used AUROC, we evaluated our model and various strategies using ARPRC ^36,37^. In the Precision-Recall space, the Precision and Recall are defined as

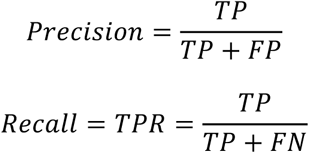

The Precision is very sensitive to FP when the number of TP is relatively small. Therefore, the AUPRC metric is able to distinguish the performances in highly unbalanced data such as the annotations of sleep arousals.

### Convolutional neural network architectures

The classic U-Net architecture was adapted in DeepSleep. The original U-Net is a 2D convolutional neural network designed for 2D image segmentation ^24^. We transformed the structure into 1D for the time-series sleep records and largely increased the number of convolutional layers from the original 18 to 35 for extracting the information at different scales. Similar to U-Net, we had convolution, max pooling and concatenation layers. The kernel size of 7 was used in the convolution operation and increasing the kernel size didn’t significantly change the performance. The nonlinear activation after each convolution operation is a Rectified Linear Unit (ReLU) defined as

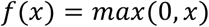

where x is the input to a neuron and f(x) is the output. Only positive values active a neuron and ReLU allows for fast and effective training of neural networks compared to other complex activation functions. In addition, batch normalization was used after each convolutional layer. In the final output layer, we used the sigmoid activation unit defined as

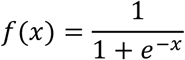

where x is the input to a neuron and f(x) is the output. During the training process, the Adam optimizer was used with the learning rate of 1e-4 and the decay rate of 1e-5.

Other network structures were also tested, including Long Short-Term Memory (LSTM) and Gated Recurrent Unit (GRU). They have similar performances. Therefore, we kept the U-Net based structure.

### Training Losses

The cross entropy loss, or log loss, was used for model training in DeepSleep. The cross entropy loss is defined as

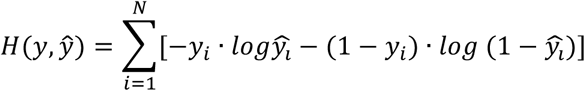

where *y*_*i*_ is the gold standard label of sleep=0 or arousal=1 at time point i, *ŷ*_*l*_ is the prediction value at time point i, N is the total number of time points, *y* is the vector of the gold standard labels and *ŷ* is the vector of predictions. Ideally, an “AUPRC loss” should be used for optimizing the prediction AUPRC. However, the “AUPRC loss” doesn’t exist because the AUPRC function is not mathematically differentiable, which is required in the neural network model training through the back-propagation algorithm ^38^. Therefore, we need to use cross-entropy loss to approximate the “AUPRC loss”. Another option is using the Sorensen-dice coefficient defined as

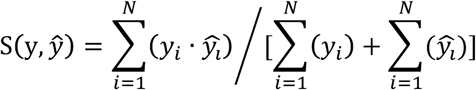

where *y*_*i*_ is the gold standard label of sleep=0 or arousal=1 at time point i, *ŷ*_*l*_ is the prediction value at time point i, N is the total number of time points, *y* is the vector of the gold standard labels and *ŷ* is the vector of predictions. We have tested the cross-entropy loss, the Sorensen dice loss and combining these two losses. Using the cross-entropy loss achieved the best performance in DeepSleep.

### Overall AUPRC and AUROC

The overall AUPRC, or the gross AUPRC, is defined as

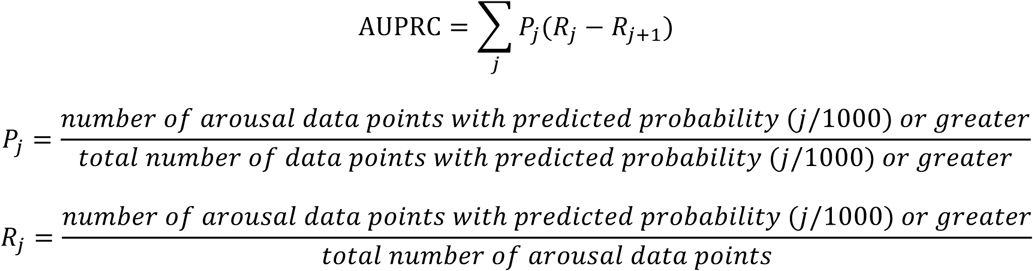

where the Precision (*P*_*j*_) and Recall (*R*_*j*_) were calculated at each cutoff j and j = 0, 0.001, 0.002, …, 0.998, 0.999, 1. For a test dataset of multiple sleep records, this overall AUPRC is similar to the “weighted AUPRC”, which is different from simply averaging the AUPRC values of all test records. This is because the overall AUPRC considers the length of each record and longer records contributing more to the overall AUPRC, resulting in a more accurate performance description of a model. The overall AUPRC was also used as the primary scoring metric in the 2018 PhysioNet Challenge. The overall AUROC was defined in a similar way as the overall AUPRC.

### Validation on the SHHS datasets

The large publicly available Sleep Heart Health Study (SHHS) dataset contains 6,441 individuals in SHHS visit 1 (SHHS1) and 3,295 individuals in SHHS visit 2 (SHHS2). The SHHS1 dataset was collected between 1995 and 1998, whereas the SHHS2 dataset was collected between 2001 and 2003. Since the recording montages were different among the PhysioNet, SHHS1, and SHHS2 datasets, the channels of polysomnograms were also different. For the SHHS1 and SHHS2 datasets, we only used a subset of 7 channels (SaO2, EEG-C3/A2, EEG-C4/A1, EOG-L, ECG, EMG, and Airflow), which were shared among these three datasets. In addition, the major signal sampling rates in the PhysioNet, SHHS1, and SHHS2 were 200Hz, 125Hz, and 250Hz respectively. We down-sample the signals to the same 25Hz by averaging successive time points. Quantile normalization was used to address the potential cohort and batch effect. For both SHHS1 and SHHS2, we randomly selected 1,000 recordings, which was comparable to the number of recordings (n=994) in the PhysioNet training dataset. Then we applied DeepSleep pipeline to train, validate and test models on SHHS1 and SHHS2 datasets individually.

## Data availability

The datasets used in this study are publicly available at the 2018 PhysioNet Challenge website and the Sleep Heart Health Study website:

https://physionet.org/physiobank/database/challenge/2018/

https://sleepdata.org/datasets/shhs

## Code availability

The code of DeepSleep is available at: https://github.com/GuanLab/DeepSleep

## Author contributions

YG and HL conceived and designed the winning algorithm in the 2018 PhysioNet Challenge. HY and YG implemented the code of various neural network structures and augmentation strategies. HY performed post-challenge analyses. All authors contributed to the writing of the manuscript and approved the final manuscript.

## Acknowledgements

This work is supported by NSF-US14-PAF07599 (CAREER: On-line Service for Predicting Protein Phosphorylation Dynamics Under Unseen Perturbations NSF), AWD007950 (Digital Biomarkers in Voices for Parkinson’s Disease American Parkinson’s Disease Association), University of Michigan O’Brien Kidney Translational Core Center, 19AMTG34850176 (American Heart Association and Amazon Web Services3.0 Data Grant Portfolio: Artificial Intelligence and Machine Learning Training Grants), and Michael J. Fox Foundation #17373. We thank the GPU donation from Nvidia.

## Supplementary Materials

### The system configuration to test DeepSleep runtimes

#### CPU

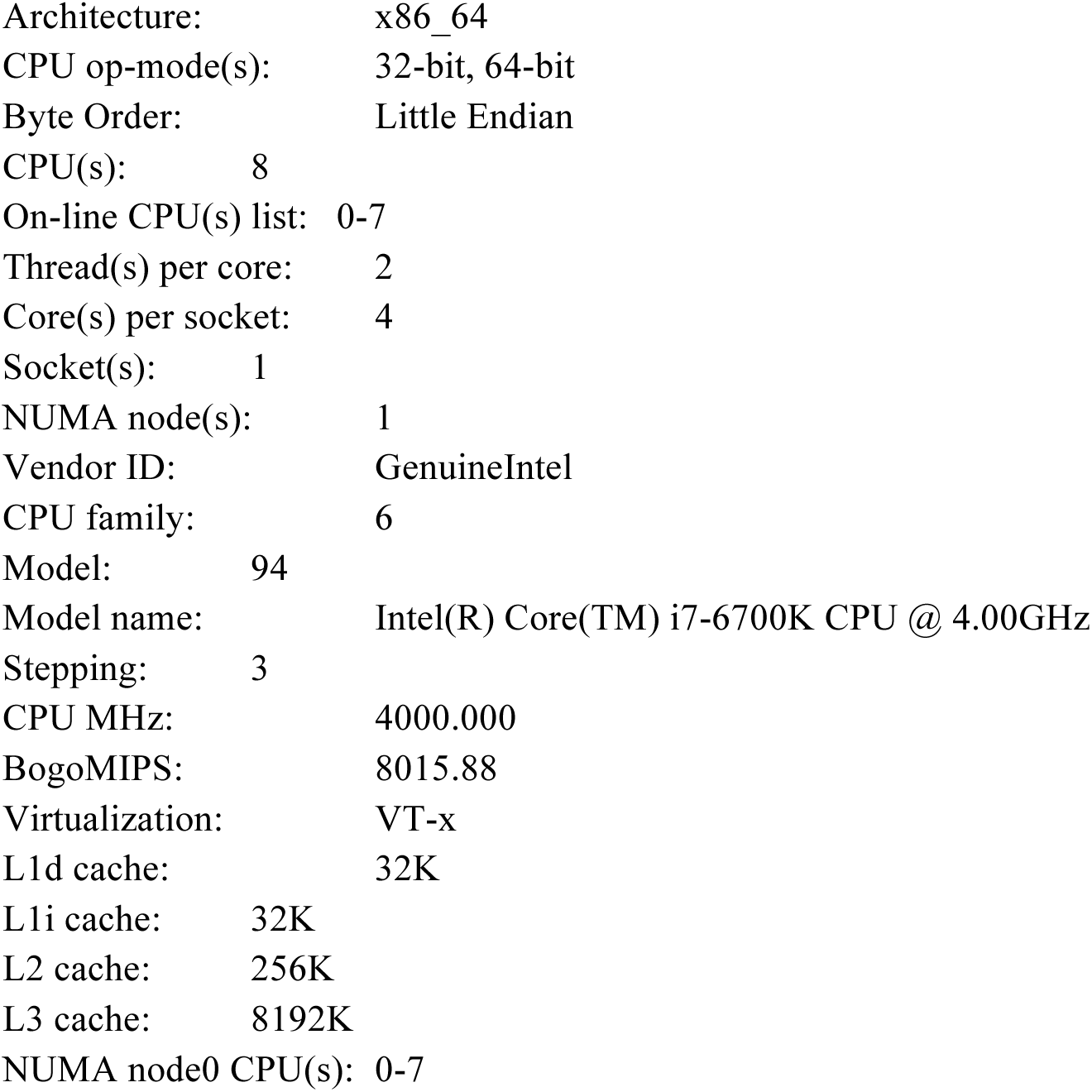

#### GPU

NVIDIA GeForce GTX TITAN X

#### Memory

31GB in total

#### System

Linux version 4.4.16-1.el7.elrepo.x86_64 (mockbuild@Build64R7) (gcc version 4.8.5 20150623 (Red Hat 4.8.5-4) (GCC)) #1 SMP Wed Jul 27 15:27:40 EDT 2016

**Fig. S1.**
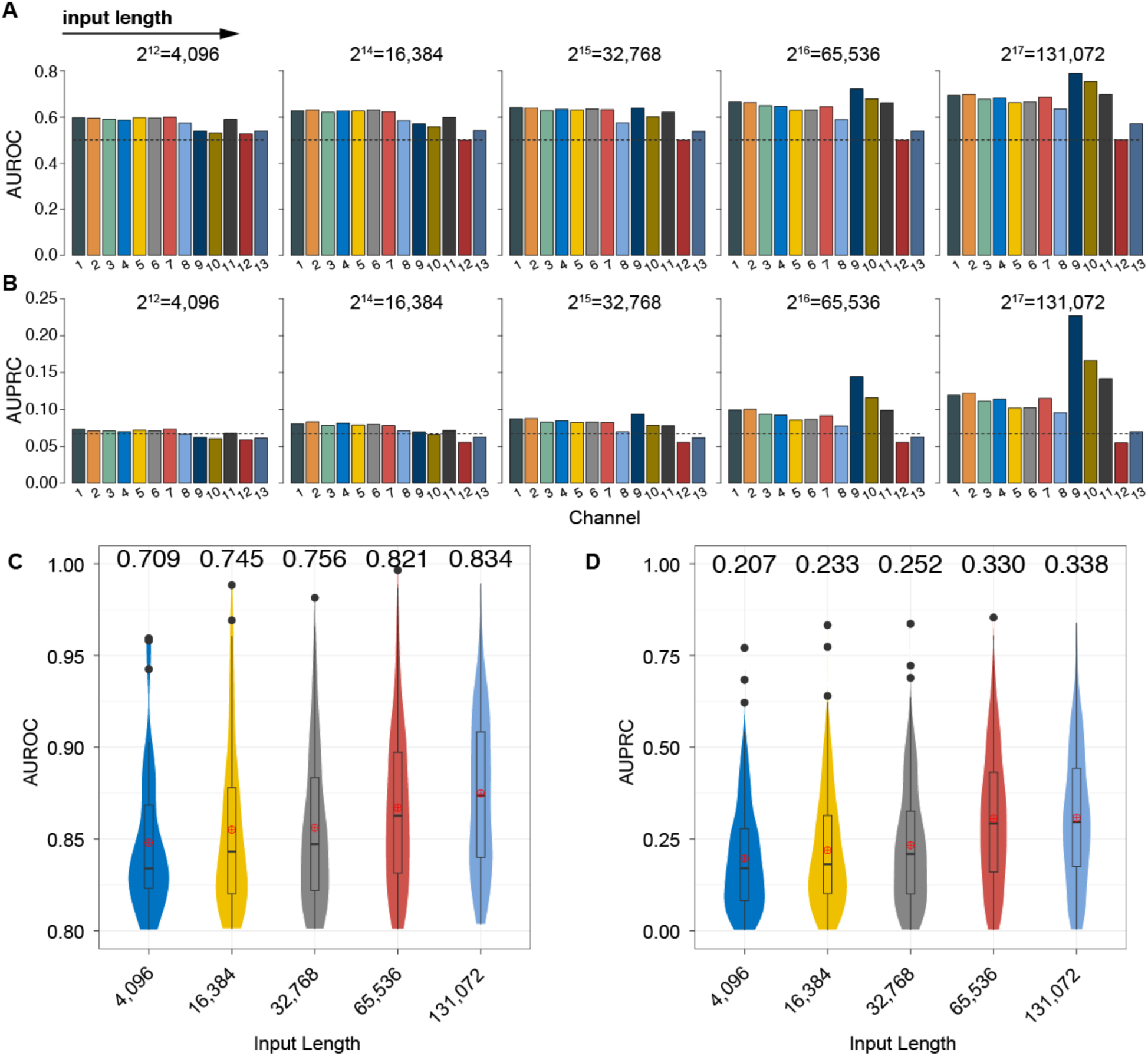
The prediction performances of models using various lengths of polysomnographic recordings as input. The (**A**) AUROCs and (**B**) AUPRCs of models using different lengths of polysomnographic recordings as input. From left to right, the length of input gradually increases from 4,096 (about 20 seconds) to 131,072 (about 11 minutes). Each color represents a model using one of the 13 polysomnographic signals. These signals correspond to the 13 channels from top to bottom in **Fig. 1 - “Data”**: 1. F3-M2; 2. F4-M1; 3. C3-M2; 4. C4-M1; 5. O1-M2; 6. O2-M1; 7. E1-M2; 8. Chin; 9. ABD; 10. Chest; 11. Airflow; 12. SaO_2_; 13. ECG. The dashed lines represent the baseline of random predictions in the AUROC space (baseline=0.500) and the AUPRC space (baseline=0.072). In contrast to (**A**) and (**B**) where a single channel was used as input, all 13 channels were used together as input features in (**C**) and (**D**). Longer input lengths achieved higher AUPRCs and AUROCs. The value above each violin is the overall AUPRC/AUROC, which is different from the simple mean or median value. The overall AUPRC/AUROC considers the length of each record and longer records contribute more to the overall AUPRC/AUROC (see details in **Methods - Overall AUPRC and AUROC**).

**Fig. S2.**
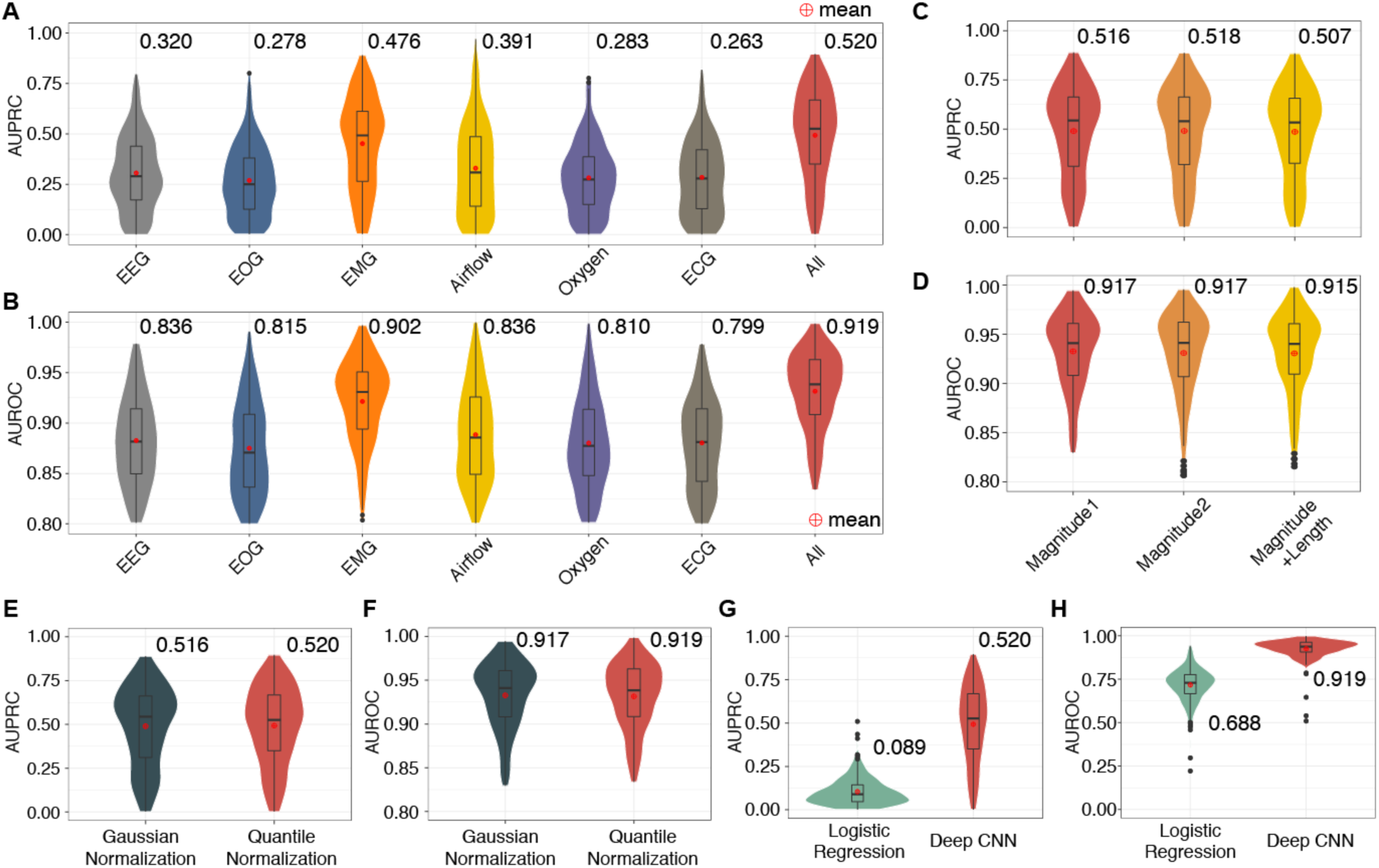
The performance comparison of models using different types of polysomnographic signals, augmentation strategies, normalization methods. From left to right, the first six categories are EEG (channel 1-6), EOG (channel 7), EMG (channel 8-10), Airflow (channel 11), saturation of Oxygen (channel 12) and ECG (channel 13). The last one, “All”, represents the model using all these 13 channels as input. The prediction (**A**) AUPRCs and (**B**) AUROCs of models using different types of signals are shown in different colors. Of note, the model “All” using all 13 polysomnographic signals achieved the best performance. We further compared the prediction (**C**) AUPRCs and (**D**) AUROCs of different data augmentation strategies are. The “Magnitude 1” strategy means that each training record was multiplied by a random number between 0.90 and 1.15, to simulate the fluctuation of the measurement in real life. The “Magnitude 2” strategy was the same as “Magnitude 1”, except for the range of the random number becomes wider, between 0.80 and 1.25. These two strategies had almost the same performance. The last “Magnitude+Length” strategy was built on top of “Magnitude 1”, in which we further extended or shrunk the record along the time axis by a random number between 0.90 and 1.15. This strategy decreased the performance and was not used in the final model training. In addition, the prediction (**E**) AUPRCs and (**F**) AUROCs of the Gaussian normalization and the quantile normalization were compared. In the Gaussian normalization, we first subtracted the average value of a signal then divided the signal by the standard deviation for each sleep record. In the quantile normalization, we first calculated the average of all training records as the reference record. Then for each record, we quantile normalized it to the reference record. The quantile normalization had better performance. We also compared the prediction (**G**) AUPRCs and (**H**) AUROCs of deep convolutional neural network (CNN) and logistic regression. Clearly, the deep CNN had much higher performance in terms of both AUPRC and AUROC. The value above each violin is the overall AUPRC/AUROC, which is different from the simple mean or median value. The overall AUPRC/AUROC considers the length of each record and longer records contribute more to the overall AUPRC/AUROC (see details in **Methods - Overall AUPRC and AUROC**).

**Fig. S3.**
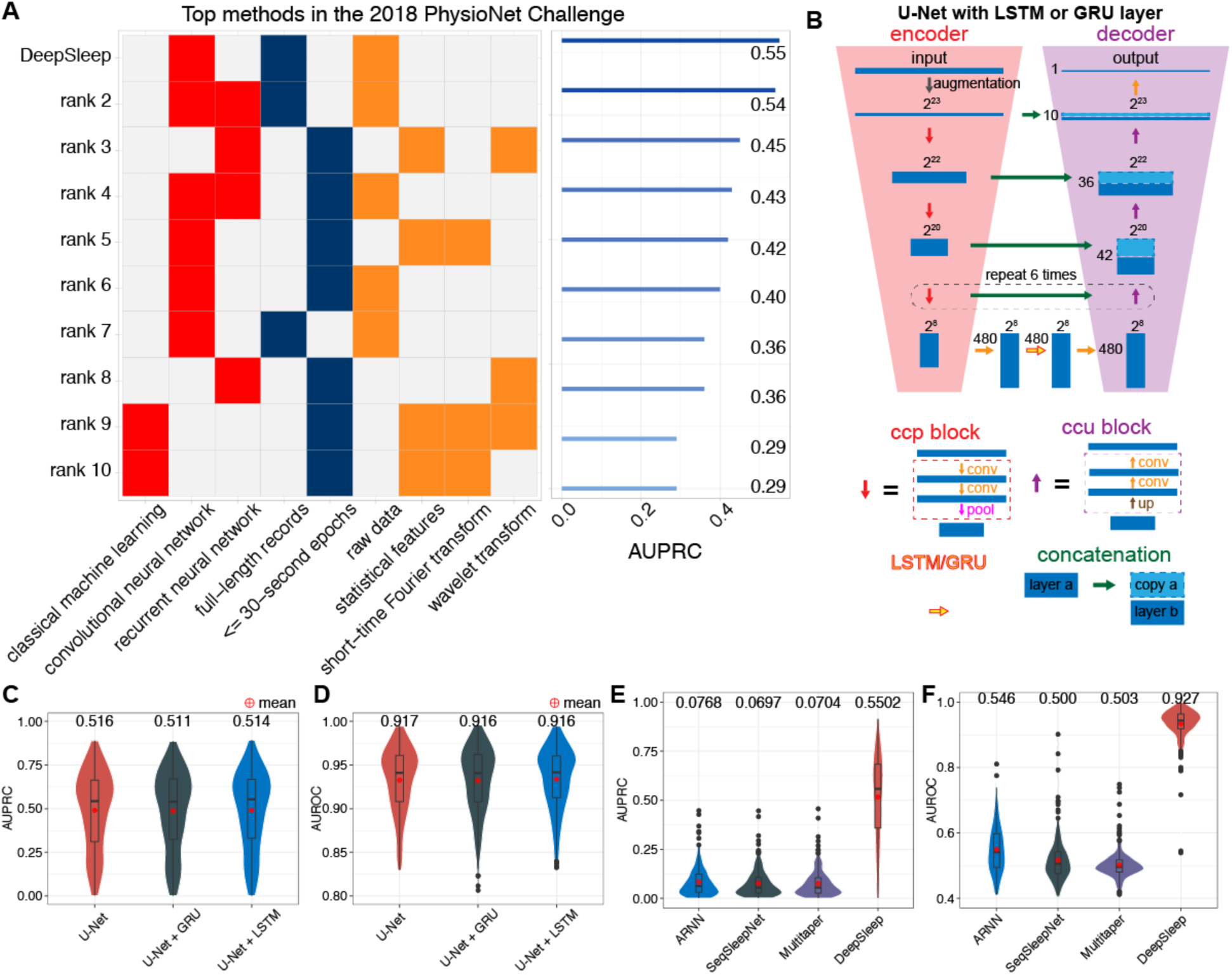
The comparison of top 10 teams in the 2018 PhysioNet Challenge, recurrent neural network, and sleep staging methods. (**A**) In the left panel, top methods, rank 2 ^1^, rank 3 ^2^, rank 4 ^3^, rank 5 ^4^, rank 6 ^5^, rank 7 ^6^, rank 8 ^7^, rank 9 ^8^, rank 10 ^9^ are compared in terms of machine learning models (red blocks), input length for models (blue blocks), and the types of input (orange blocks). In particular, the input are either raw polysomnogram data, or features extracted by statistical analysis, short-time Fourier transform, or wavelet transform. The corresponding prediction performances of these methods are shown in the right panel. We also implemented the recurrent neural network (RNN) structure by adding a recurrent unit of LSTM or GRU layer (yellow arrow with red border) at the bottom of U-Net (**B**). The arrows in different colors represent different neural network layers, blocks or operations. The prediction (**C**) AUPRCs and (**D**) AUROCs of U-Net, U-Net with GRU and U-Net with LSTM are shown in different colors. Adding the recurrent layer did not improve the performance. We used U-Net without recurrent layers as in our final model. We further compared current methods for sleep staging. The prediction (**E**) AUPRCs and (**F**) AUROCs of (a) attention recurrent neural network (ARNN) ^10^, (b) SeqSleepNet using features from short-time Fourier transform ^11,12^, (c) a method using features from Thomson’s multitaper ^13,14^, and (d) our DeepSleep approach are shown in different colors. The value above each violin is the overall AUPRC/AUROC, which is different from the simple mean or median value. The overall AUPRC/AUROC considers the length of each record and longer records contribute more to the overall AUPRC/AUROC (see details in **Methods - Overall AUPRC and AUROC**).

**Fig. S4.**
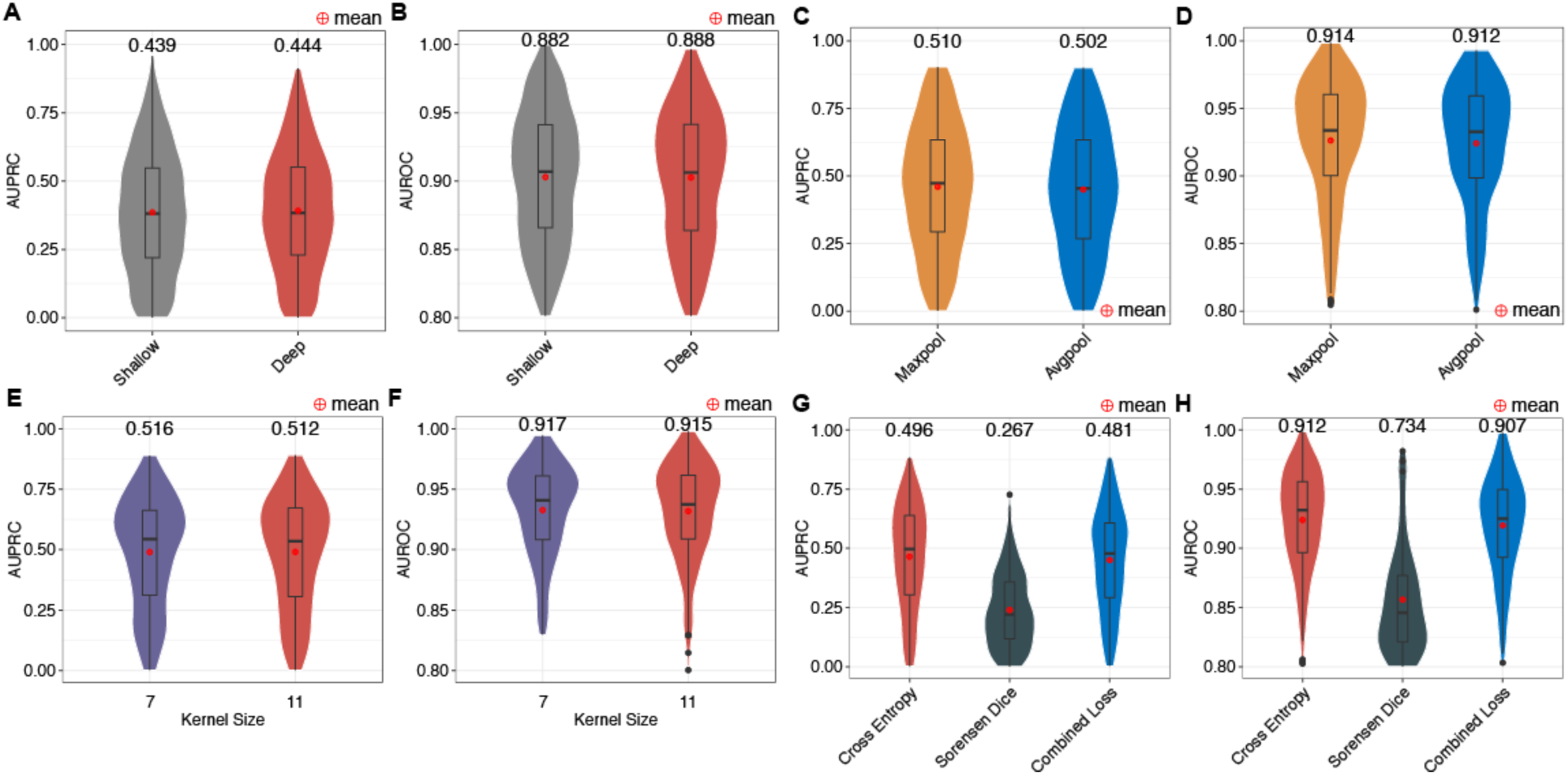
The performance comparison of U-Net with different modifications. The prediction (**A**) AUPRCs and (**B**) AUROCs of the “Shallow” and “Deep” U-Net were compared. The “Shallow” structure is only relatively shallow (4 less convolutional layers), compared with the “Deep” structure. Nevertheless, the “Shallow” U-Net already showed worse prediction performance than the “Deep” one. The prediction (**C**) AUPRCs and (**D**) AUROCs of U-Net with the kernel size of 7 and 11 in the convolutional layers were compared. Since the performances were very similar and the kernel size of 11 required more computational time and sources, we used the kernel size of 7 in our model. The prediction (**E**) AUPRCs and (**F**) AUROCs of U-Net with max-pooling or average-pooling layers are also compared. Using max-pooling layers has slightly higher performance, which was implemented in our model. The prediction (**G**) AUPRCs and (**H**) AUROCs of models trained with the cross-entropy loss, the sorensen dice loss or combining both losses were further tested. The cross-entropy loss significantly outperformed the sorensen dice loss. Even if we combined both losses, the performance was still lower. Therefore, we used the cross-entropy loss function to train our model. The value above each model is the overall AUPRC/AUROC, which is different from the simple mean or median value. The overall AUPRC/AUROC considers the length of each record and longer records contribute more to the overall AUPRC/AUROC (see details in **Methods - Overall AUPRC and AUROC**).

**Fig. S5.**
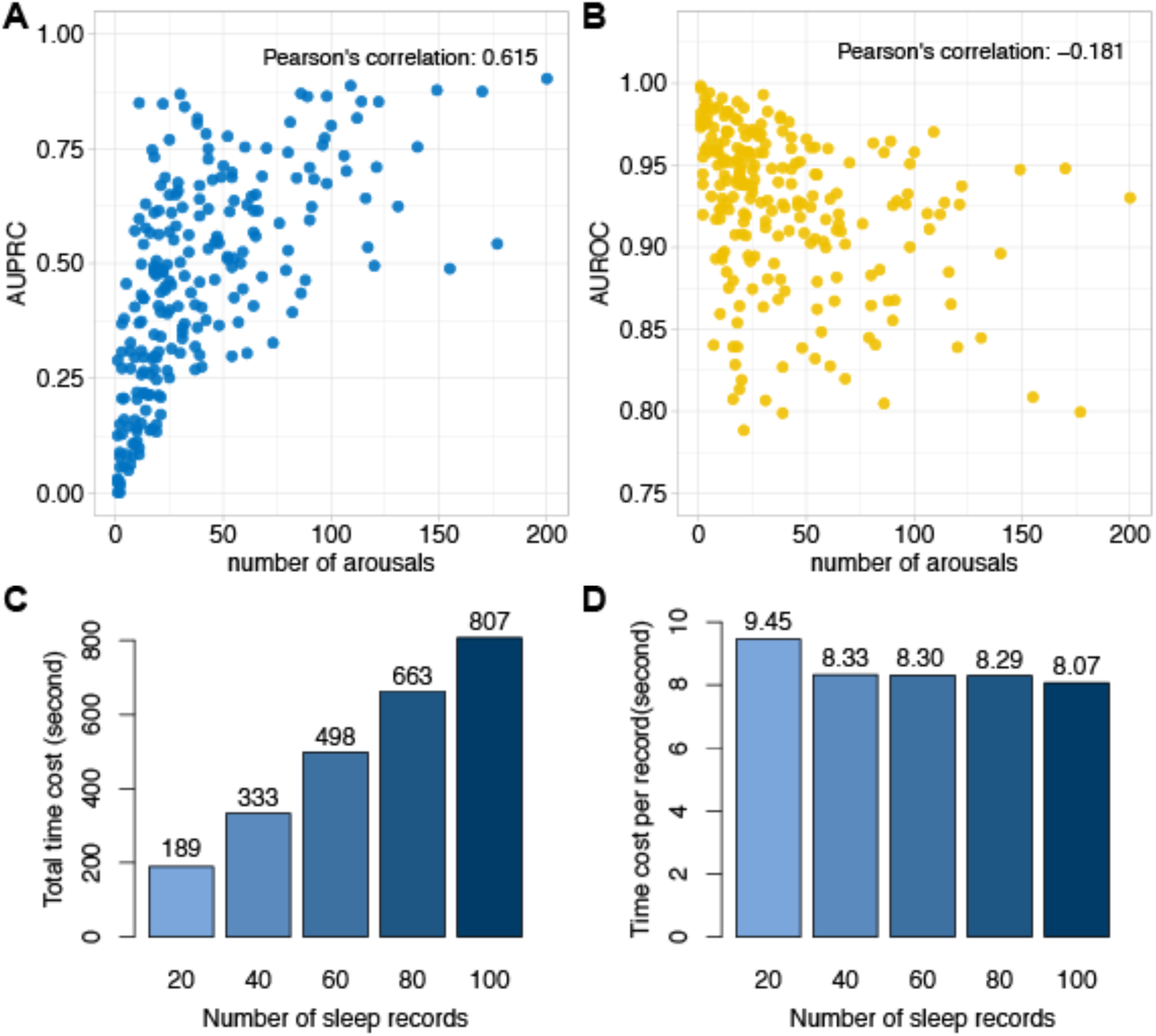
The relationship between prediction performance and the number of arousals, and the runtimes for predicting sleep arousals. The prediction (**A**) AUPRCs and (**B**) AUROCs are shown by the y-axis. Each dot represents one sleep record. The AUPRC has a medium correlation with the number of sleep arousals. The (**C**) total time cost and (**D**) average time cost per sleep record are shown in bar plots. Notably, the average runtime per sleep record is less than 10 seconds and gradually decreases as the total number of records to be analyzed increases. This results from the overhead time of loading the large neural network models before the prediction step.

**Fig. S6.**
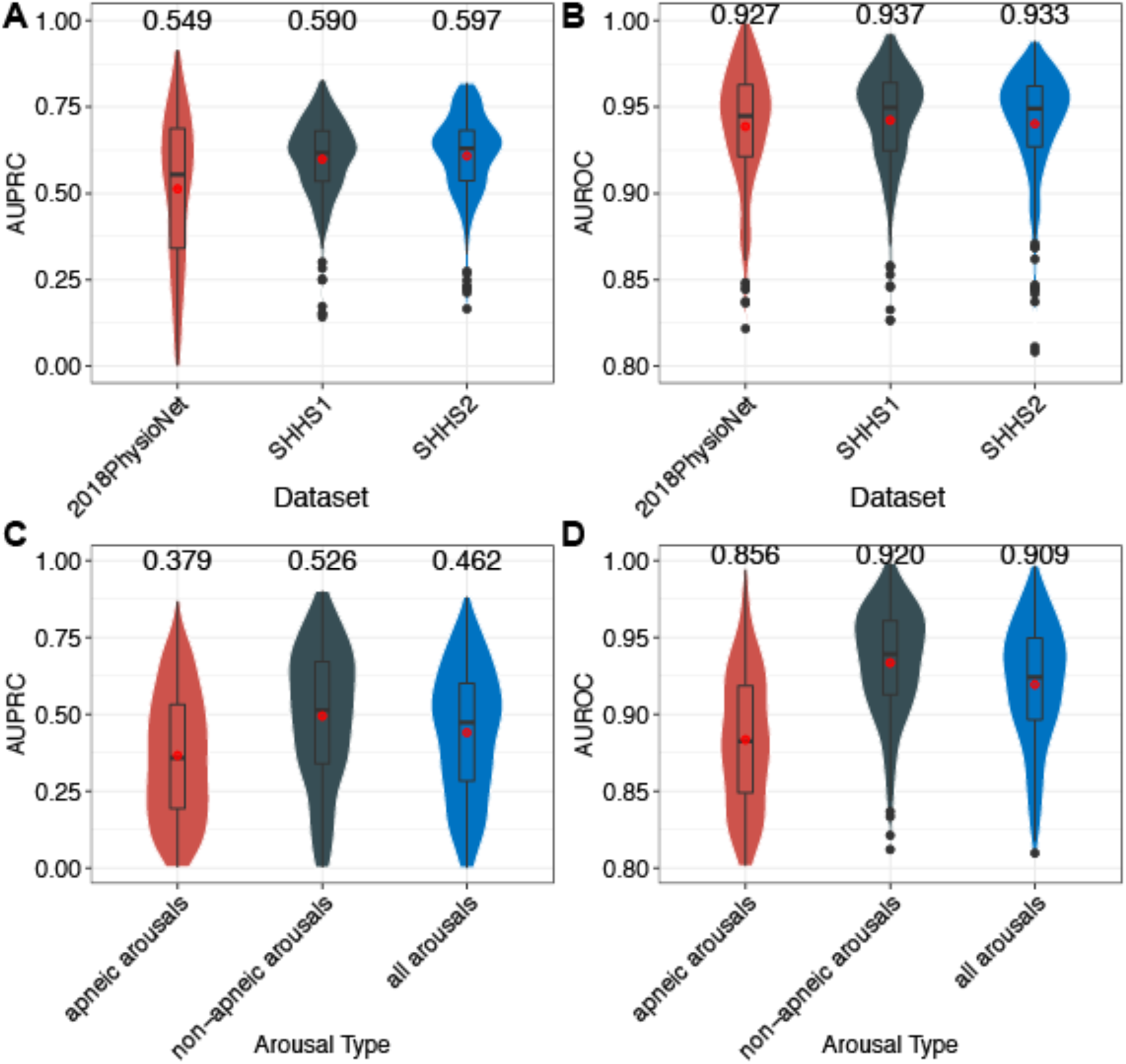
The performance comparison of DeepSleep on different datasets and different types of arousals. The prediction (**A**) AUPRCs and (**B**) AUROCs of DeepSleep on the 2018-PhysioNet, Sleep Heart Health Study visit 1 (SHHS1), and SHHS2 datasets were compared. The performance on these three datasets was comparable. We further tested the prediction (**C**) AUPRCs and (**D**) AUROCs of DeepSleep on apneic, non-apneic, and all (both apneic and non-apneic) arousals. The value above each violin is the overall AUPRC/AUROC, which is different from the simple mean or median value. The overall AUPRC/AUROC considers the length of each record and longer records contribute more to the overall AUPRC/AUROC (see details in **Methods - Overall AUPRC and AUROC**).

**Fig. S7.**
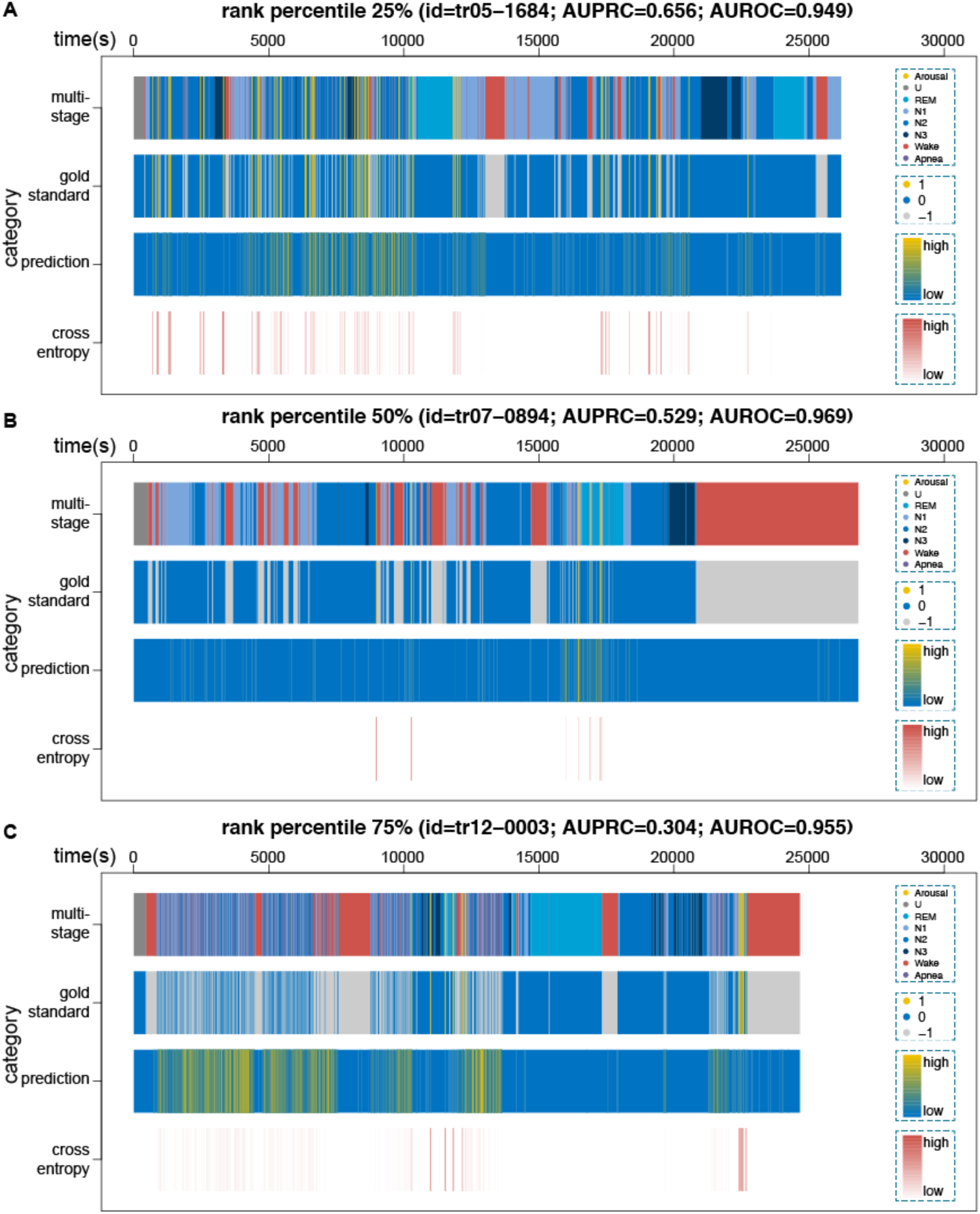
Visualization of our prediction and the gold standard annotation for three sleep records with rank percentile 25%, 50%, and 75% based on the prediction AUPRC. From top to bottom along the y-axis, the four rows correspond to the 8 annotation categories, the binary label of arousal (yellow) and sleep (blue), excluding the non-scoring regions (gray), the continuous prediction and the cross entropy loss at each data point. The sleep records in (**A**), (**B**), and (**C**) were ranked 25%, 50%, and 75% respectively among all records based on the prediction AUPRC.

**Table S1.**
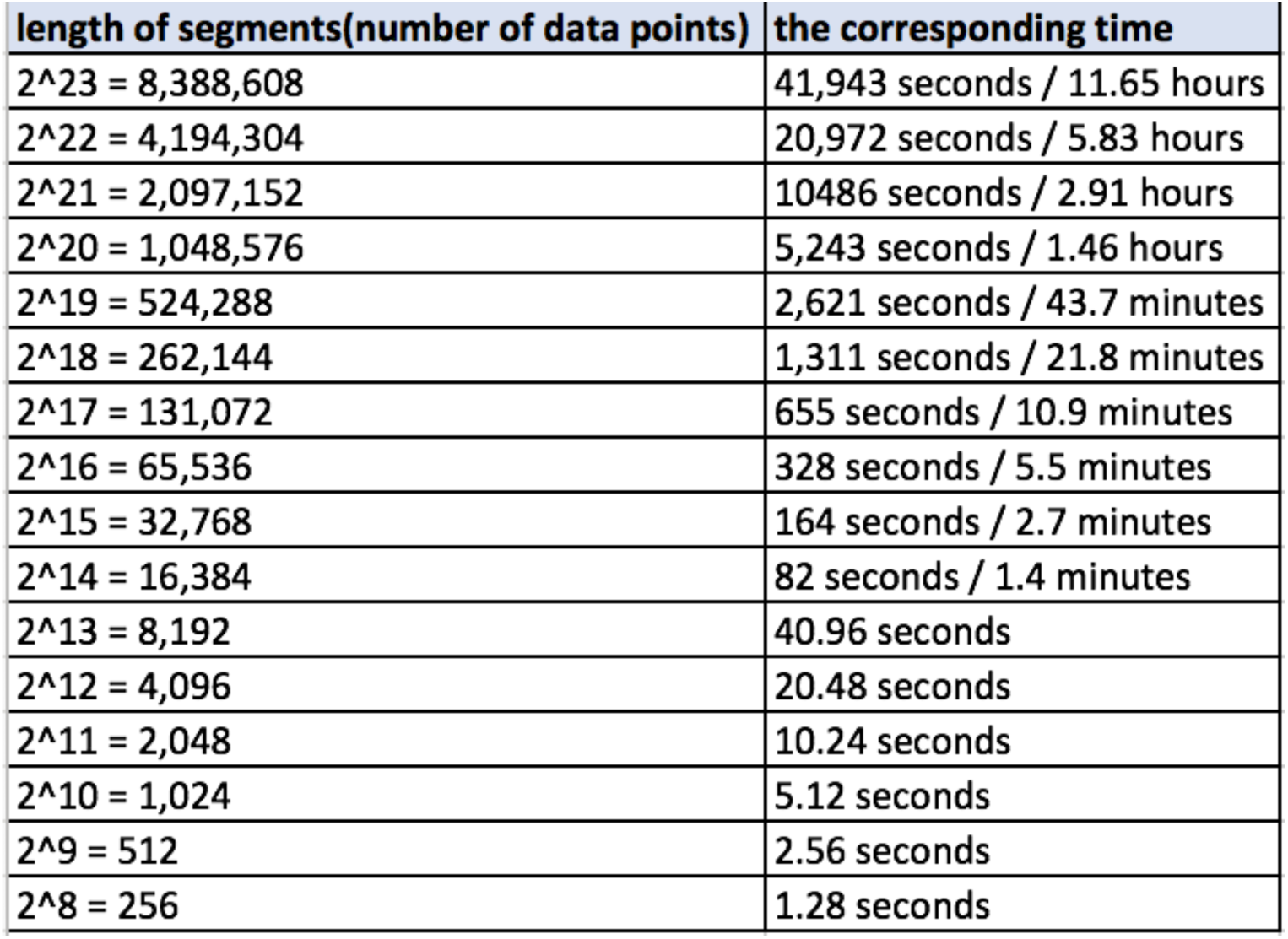
The relationship between length of segments and the corresponding time.

