## Supplemental Information for "DeepSleep: Fast and Accurate Delineation of Sleep Arousals at Millisecond Resolution by Deep Learning"

**The system configuration to test DeepSleep runtimes**

**CPU**

| | |
|---|---|
| Architecture: | x86_64 |
| CPU op-mode(s): | 32-bit, 64-bit |
| Byte Order: | Little Endian |
| CPU(s): | 8 |
| On-line CPU(s) list: | 0-7 |
| Thread(s) per core: | 2 |
| Core(s) per socket: | 4 |
| Socket(s): | 1 |
| NUMA node(s): | 1 |
| Vendor ID: | GenuineIntel |
| CPU family: | 6 |
| Model: | 94 |
| Model name: | Intel(R) Core(TM) i7-6700K CPU @ 4.00GHz |
| Stepping: | 3 |
| CPU MHz: | 4000.000 |
| BogoMIPS: | 8015.88 |
| Virtualization: | VT-x |
| L1d cache: | 32K |
| L1i cache: | 32K |
| L2 cache: | 256K |
| L3 cache: | 8192K |
| NUMA node0 CPU(s): | 0-7 |

**GPU**

NVIDIA GeForce GTX TITAN X

**Memory**

31GB in total

**System**

Linux version 4.4.16-1.el7.elrepo.x86_64 (mockbuild@Build64R7) (gcc version 4.8.5 20150623 (Red Hat 4.8.5-4) (GCC) ) #1 SMP Wed Jul 27 15:27:40 EDT 2016

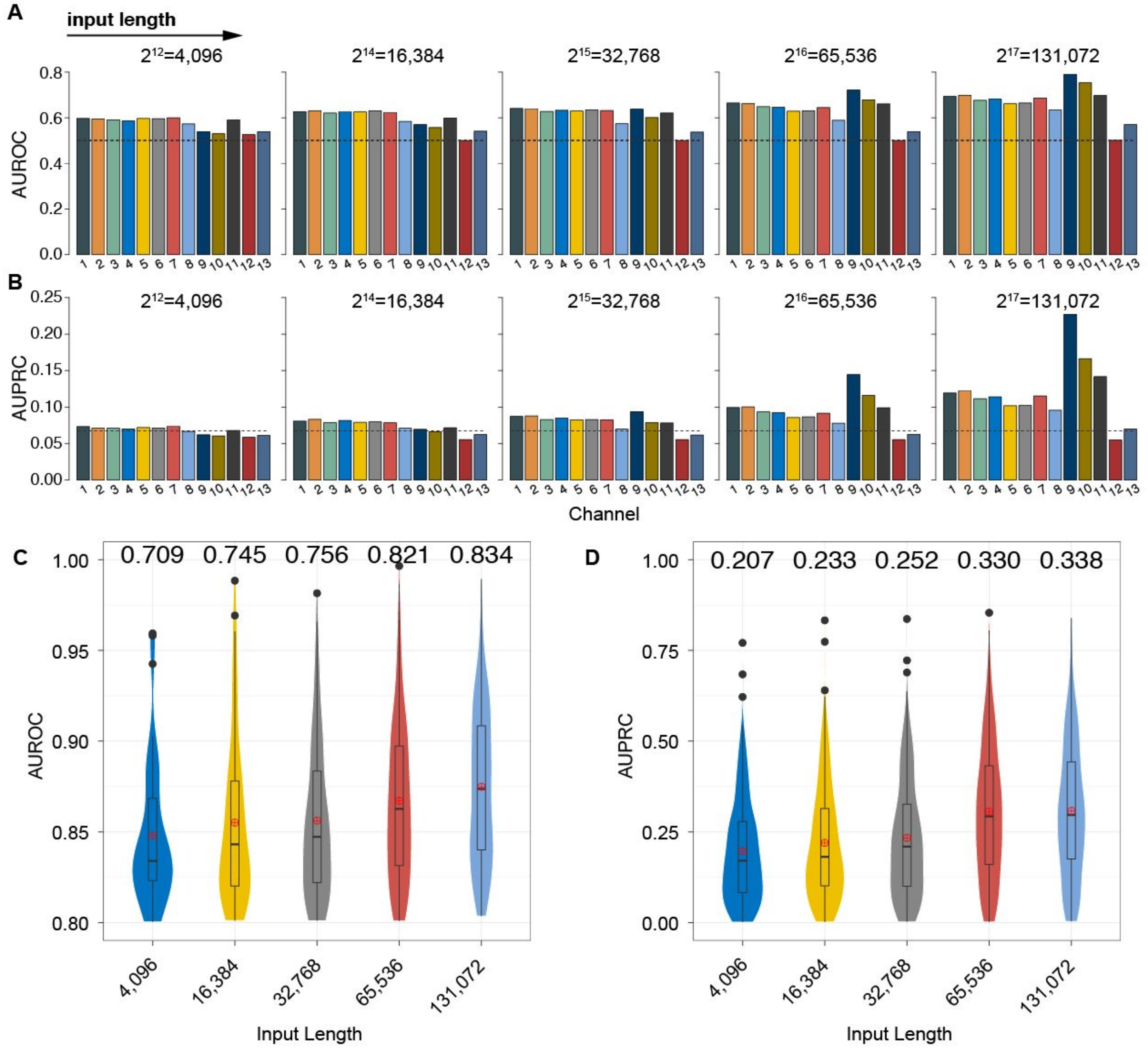


**Fig. S1. The prediction performances of models using various lengths of polysomnographic recordings as input.**

The (**A**) AUROCs and (**B**) AUPRCs of models using different lengths of polysomnographic recordings as input. From left to right, the length of input gradually increases from 4,096 (about 20 seconds) to 131,072 (about 11 minutes). Each color represents a model using one of the 13 polysomnographic signals. These signals correspond to the 13 channels from top to bottom in **Fig. 1 - "Data"**: 1. F3-M2; 2. F4-M1; 3. C3-M2; 4. C4-M1; 5. O1-M2; 6. O2-M1; 7. E1-M2; 8. Chin; 9. ABD; 10. Chest; 11. Airflow; 12. $SaO_2$; 13. ECG. The dashed lines represent the baseline of random predictions in the AUROC space (baseline=0.500) and the AUPRC space (baseline=0.072). In contrast to (**A**) and (**B**) where a single channel was used as input, all 13 channels were used together as input features in (**C**) and (**D**). Longer input lengths achieved higher AUPRCs and AUROCs. The value above each violin is the overall AUPRC/AUROC, which is different from the simple mean or median value. The overall

AUPRC/AUROC considers the length of each record and longer records contribute more to the overall AUPRC/AUROC (see details in **Methods - Overall AUPRC and AUROC**).

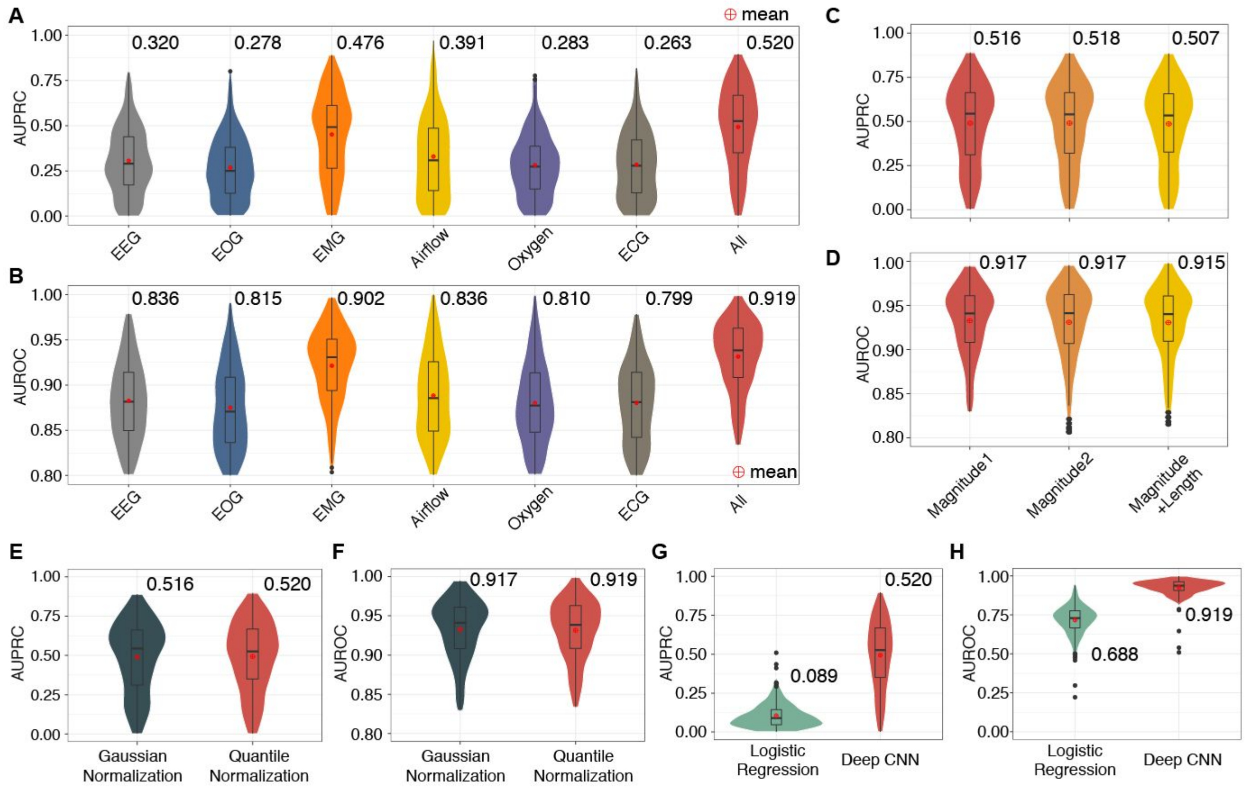


**Fig. S2. The performance comparison of models using different types of polysomnographic signals, augmentation strategies, normalization methods.**

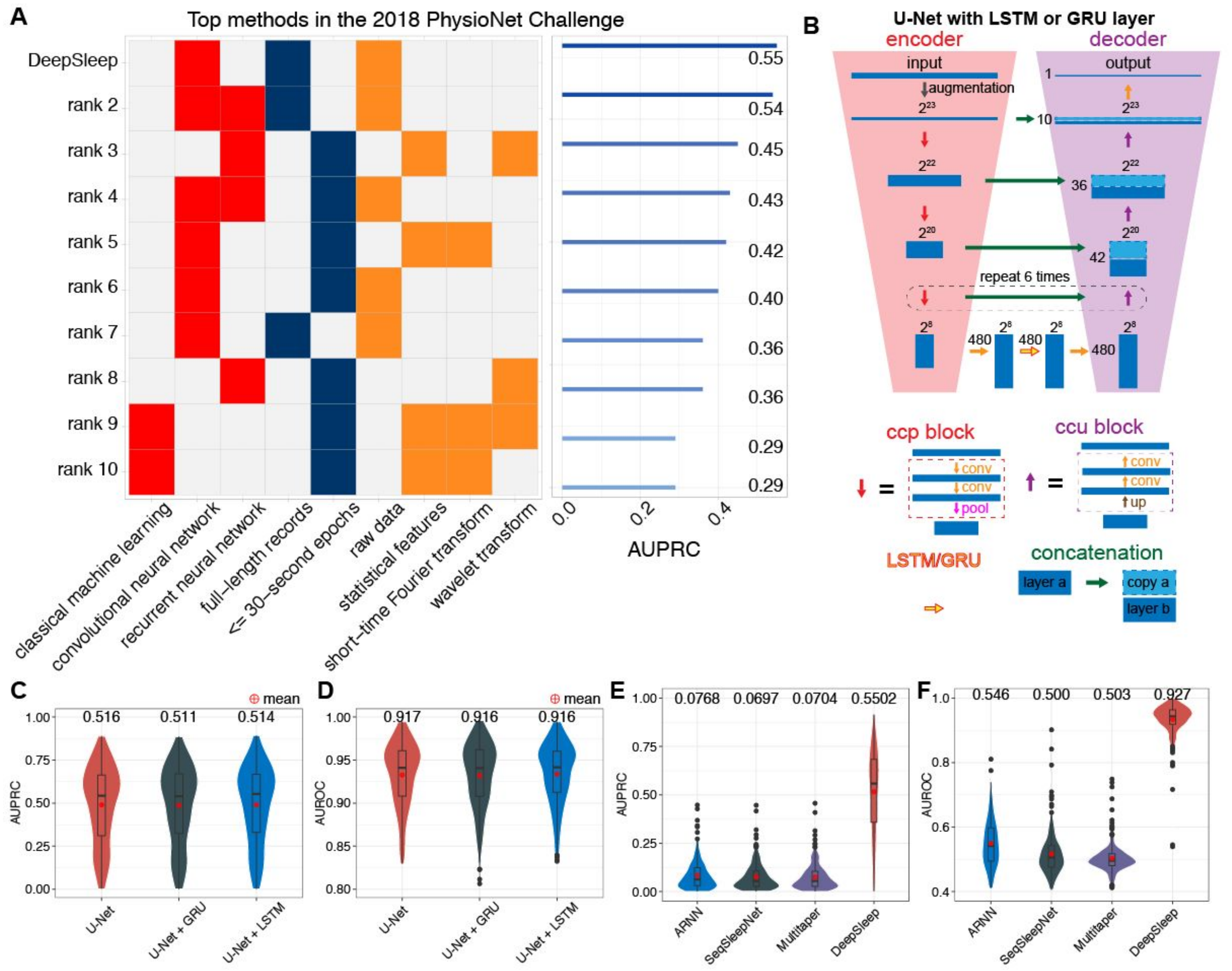


**Fig. S3. The comparison of top 10 teams in the 2018 PhysioNet Challenge, recurrent neural network, and sleep staging methods.**

(**A**) In the left panel, top methods, rank 2 [1], rank 3 [2], rank 4 [3], rank 5 [4], rank 6 [5], rank 7 [6], rank 8 [7], rank 9 [8], rank 10 [9] are compared in terms of machine learning models (red blocks), input length for models (blue blocks), and the types of input (orange blocks). In particular, the input are either raw polysomnogram data, or features extracted by statistical analysis, short-time Fourier transform, or wavelet transform. The corresponding prediction performances of these methods are shown in the right panel. We also implemented the recurrent neural network (RNN) structure by adding a recurrent unit of LSTM or GRU layer (yellow arrow with red border) at the bottom of U-Net (**B**). The arrows in different colors represent different neural network layers, blocks or operations. The prediction (**C**) AUPRCs and (**D**) AUROCs of U-Net, U-Net with GRU and U-Net with LSTM are shown in different colors. Adding the recurrent layer did not improve the performance. We used U-Net without recurrent layers as in our final model. We further compared current methods for sleep staging. The prediction (**E**) AUPRCs and (**F**) AUROCs of (a) attention recurrent neural network (ARNN) [10], (b) SeqSleepNet using features from short-time Fourier transform [11,12], (c) a method using features from Thomson's multitaper [13,14], and (d) our DeepSleep approach are shown in different colors. The value above each violin is the overall AUPRC/AUROC, which is different from the simple mean or median value. The overall AUPRC/AUROC considers the length of each record and longer records contribute more to the overall AUPRC/AUROC (see details in **Methods - Overall AUPRC and AUROC**).

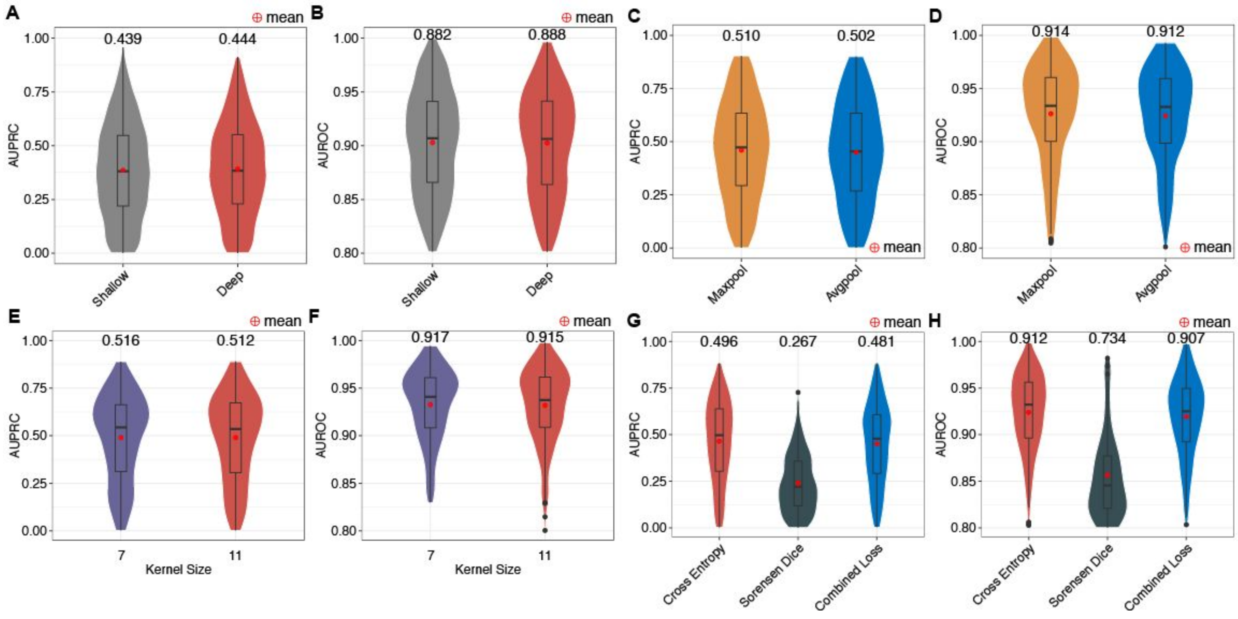


**Fig. S4. The performance comparison of U-Net with different modifications.**

The prediction (**A**) AUPRCs and (**B**) AUROCs of the "Shallow" and "Deep" U-Net were compared. The "Shallow" structure is only relatively shallow (4 less convolutional layers), compared with the "Deep" structure. Nevertheless, the "Shallow" U-Net already showed worse prediction performance than the "Deep" one. The prediction (**C**) AUPRCs and (**D**) AUROCs of U-Net with the kernel size of 7 and 11 in the convolutional layers were compared. Since the performances were very similar and the kernel size of 11 required more computational time and sources, we used the kernel size of 7 in our model. The prediction (**E**) AUPRCs and (**F**) AUROCs of U-Net with max-pooling or average-pooling layers are also compared. Using max-pooling layers has slightly higher performance, which was implemented in our model. The prediction (**G**) AUPRCs and (**H**) AUROCs of models trained with the cross-entropy loss, the sorensen dice loss or combining both losses were further tested. The cross-entropy loss significantly outperformed the sorensen dice loss. Even if we combined both losses, the performance was still lower.

Therefore, we used the cross-entropy loss function to train our model. The value above each model is the overall AUPRC/AUROC, which is different from the simple mean or median value. The overall AUPRC/AUROC considers the length of each record and longer records contribute more to the overall AUPRC/AUROC (see details in **Methods - Overall AUPRC and AUROC**).

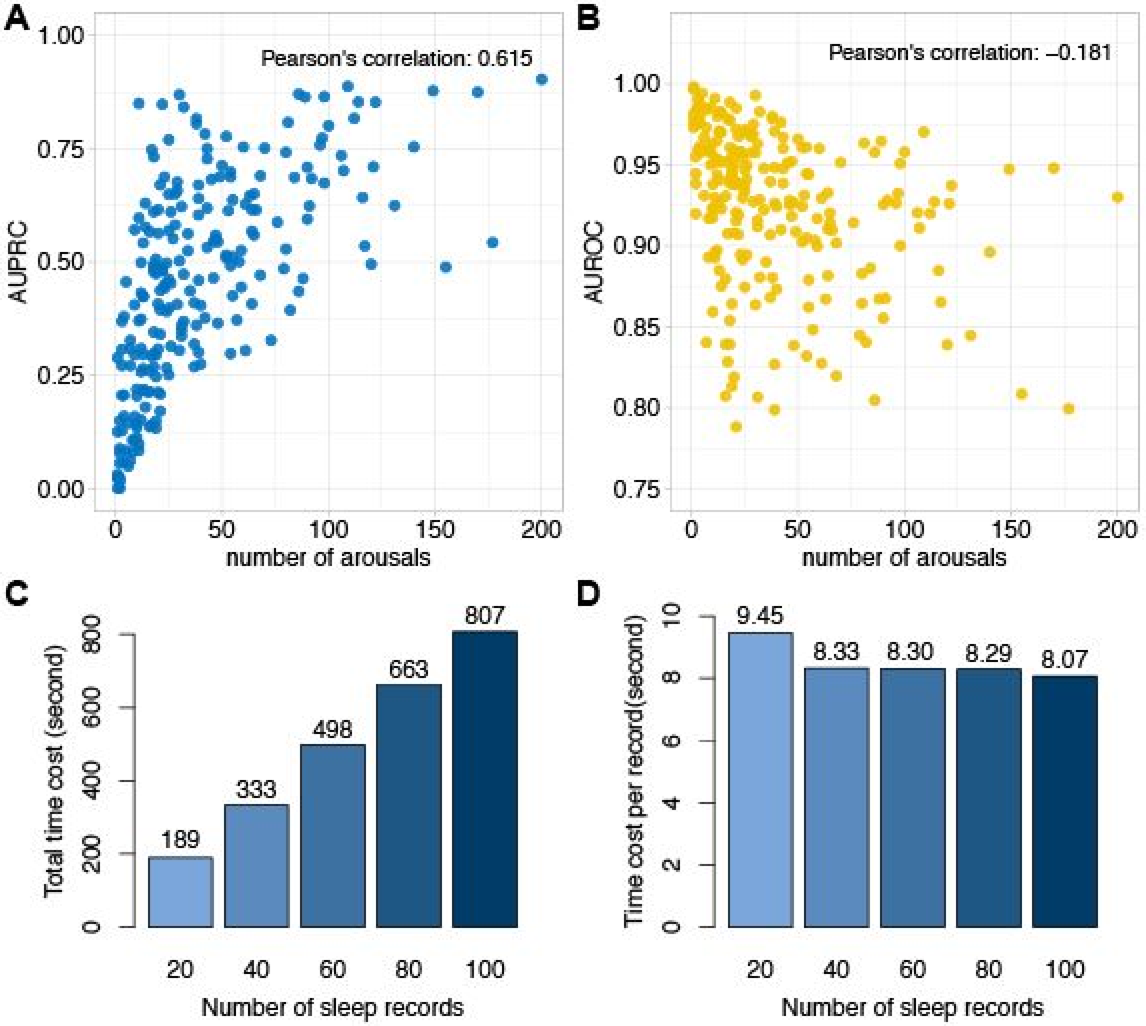


**Fig. S5. The relationship between prediction performance and the number of arousals, and the runtimes for predicting sleep arousals.**

The prediction (**A**) AUPRCs and (**B**) AUROCs are shown by the y-axis. Each dot represents one sleep record. The AUPRC has a medium correlation with the number of sleep arousals. The (**C**) total time cost and (**D**) average time cost per sleep record are shown in bar plots. Notably, the average runtime per sleep record is less than 10 seconds and gradually decreases as the total number of records to be analyzed increases. This results from the overhead time of loading the large neural network models before the prediction step.

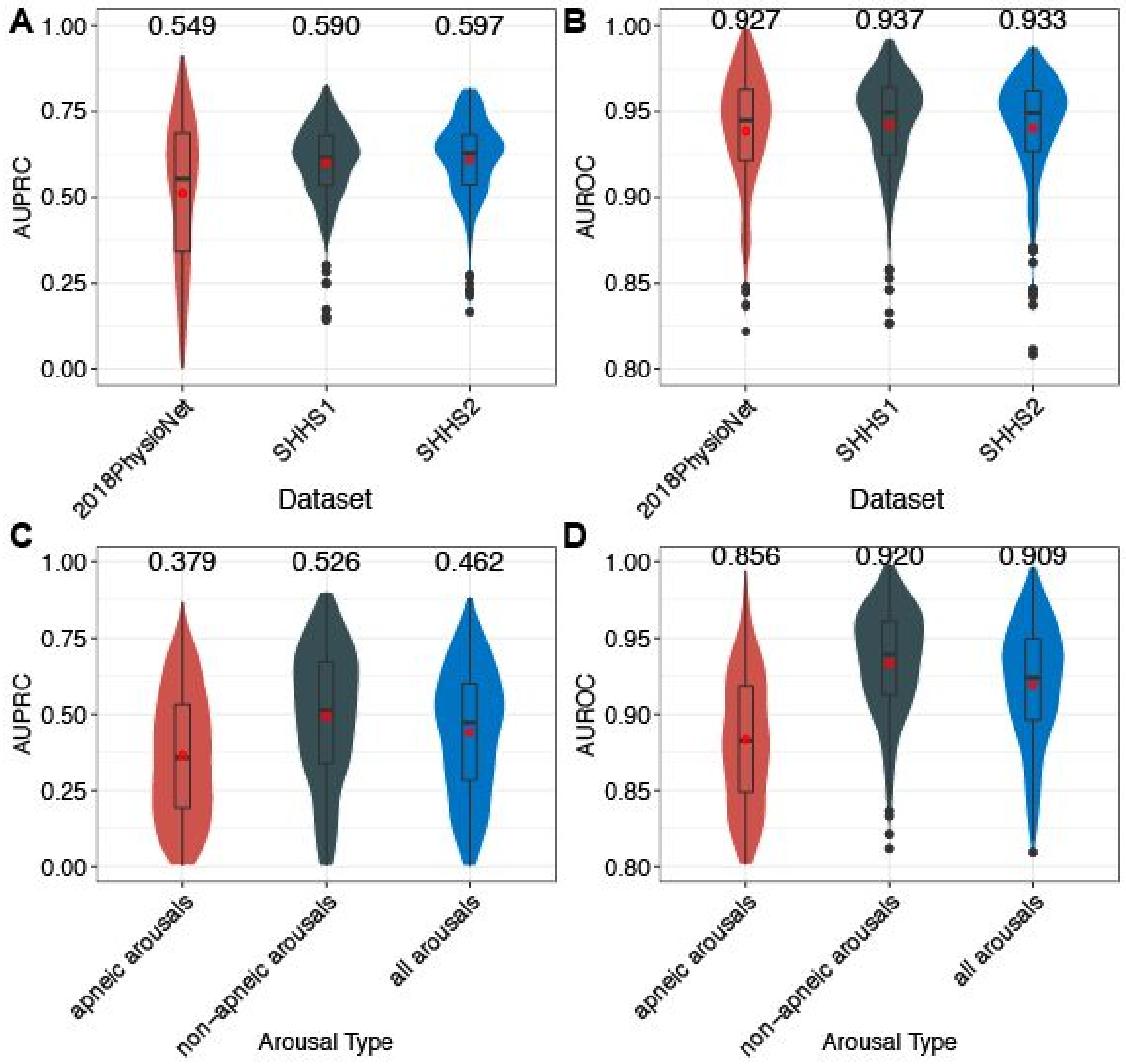


**Fig. S6. The performance comparison of DeepSleep on different datasets and different types of arousals**

The prediction (**A**) AUPRCs and (**B**) AUROCs of DeepSleep on the 2018-PhysioNet, Sleep Heart Health Study visit 1 (SHHS1), and SHHS2 datasets were compared. The performance on these three datasets was comparable. We further tested the prediction (**C**) AUPRCs and (**D**) AUROCs of DeepSleep on apneic, non-apneic, and all (both apneic and non-apneic) arousals. The value above each violin is the overall AUPRC/AUROC, which is different from the simple mean or median value. The overall AUPRC/AUROC considers the length of each record and longer records contribute more to the overall AUPRC/AUROC (see details in **Methods - Overall AUPRC and AUROC**).

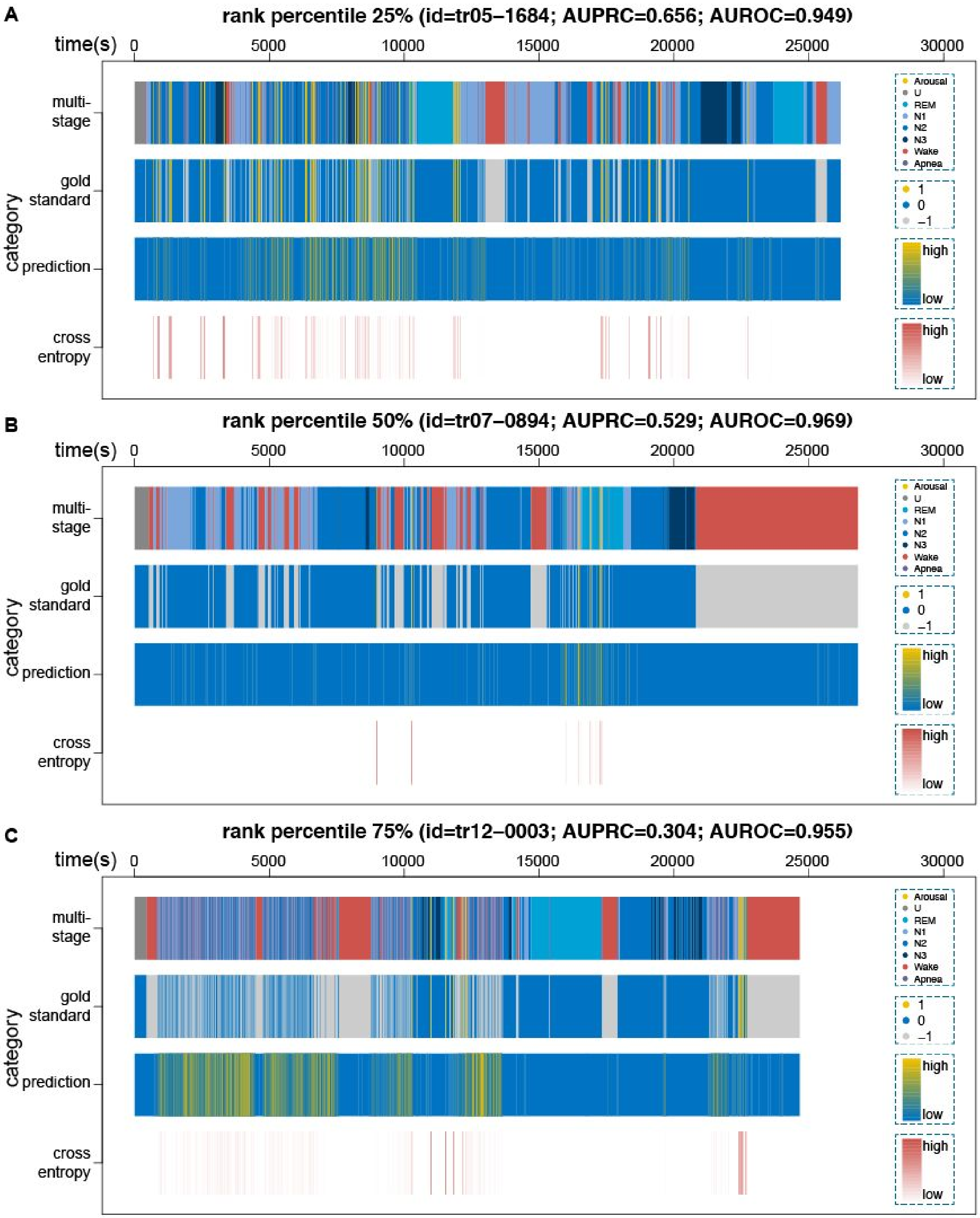
A
rank percentile 25% (id=tr05–1684; AUPRC=0.656; AUROC=0.949)
B
rank percentile 50% (id=tr07–0894; AUPRC=0.529; AUROC=0.969)
C
rank percentile 75% (id=tr12–0003; AUPRC=0.304; AUROC=0.955)
time(s)
0
5000
10000
15000
20000
25000
30000
category
multi-stage
gold standard
prediction
cross entropy
Arousal
U
REM
N1
N2
N3
Wake
Apnea
1
0
–1
high
low

**Fig. S7. Visualization of our prediction and the gold standard annotation for three sleep records with rank percentile 25%, 50%, and 75% based on the prediction AUPRC.**

From top to bottom along the y-axis, the four rows correspond to the 8 annotation categories, the binary label of arousal (yellow) and sleep (blue), excluding the non-scoring regions (gray), the continuous prediction and the cross entropy loss at each data point. The sleep records in (**A**), (**B**), and (**C**) were ranked 25%, 50%, and 75% respectively among all records based on the prediction AUPRC.

**Table S1. The relationship between length of segments and the corresponding time.**

| length of segments(number of data points) | the corresponding time |
|---|---|
| 2^23 = 8,388,608 | 41,943 seconds / 11.65 hours |
| 2^22 = 4,194,304 | 20,972 seconds / 5.83 hours |
| 2^21 = 2,097,152 | 10486 seconds / 2.91 hours |
| 2^20 = 1,048,576 | 5,243 seconds / 1.46 hours |
| 2^19 = 524,288 | 2,621 seconds / 43.7 minutes |
| 2^18 = 262,144 | 1,311 seconds / 21.8 minutes |
| 2^17 = 131,072 | 655 seconds / 10.9 minutes |
| 2^16 = 65,536 | 328 seconds / 5.5 minutes |
| 2^15 = 32,768 | 164 seconds / 2.7 minutes |
| 2^14 = 16,384 | 82 seconds / 1.4 minutes |
| 2^13 = 8,192 | 40.96 seconds |
| 2^12 = 4,096 | 20.48 seconds |
| 2^11 = 2,048 | 10.24 seconds |
| 2^10 = 1,024 | 5.12 seconds |
| 2^9 = 512 | 2.56 seconds |
| 2^8 = 256 | 1.28 seconds |

2. Már Þráinsson, H. *et al.* Automatic Detection of Target Regions of Respiratory Effort-Related

Arousals Using Recurrent Neural Networks. in *2018 Computing in Cardiology Conference (CinC)* **45**, (Computing in Cardiology, 2018).

3. He, R. *et al.* Identification of Arousals With Deep Neural Networks Using Different Physiological Signals. in *2018 Computing in Cardiology Conference (CinC)* **45**, (Computing in Cardiology, 2018).
4. Varga, B., Görög, M. & Hajas, P. Using Auxiliary Loss to Improve Sleep Arousal Detection With Neural Network. in *2018 Computing in Cardiology Conference (CinC)* **45**, (Computing in Cardiology, 2018).
5. Patane, A., Ghiasi, S., Pasquale Scilingo, E. & Kwiatkowska, M. Automated Recognition of Sleep Arousal Using Multimodal and Personalized Deep Ensembles of Neural Networks. in *2018 Computing in Cardiology Conference (CinC)* **45**, (Computing in Cardiology, 2018).
6. Miller, D., Ward, A. & Bambos, N. Automatic Sleep Arousal Identification From Physiological Waveforms Using Deep Learning. in *2018 Computing in Cardiology Conference (CinC)* **45**, (Computing in Cardiology, 2018).
7. Warrick, P. & Nabhan Homsi, M. Sleep Arousal Detection From Polysomnography Using the Scattering Transform and Recurrent Neural Networks. in *2018 Computing in Cardiology Conference (CinC)* **45**, (Computing in Cardiology, 2018).
8. Bhattacharjee, T. *et al.* SleepTight: Identifying Sleep Arousals Using Inter and Intra-Relation of Multimodal Signals. in *2018 Computing in Cardiology Conference (CinC)* **45**, (Computing in Cardiology, 2018).
9. Szalma, J., Bánhalmi, A. & Bilicki, V. Detection of Respiratory Effort-Related Arousals Using a Hidden Markov Model and Random Decision Forest. in *2018 Computing in Cardiology Conference (CinC)* **45**, (Computing in Cardiology, 2018).
10. Phan, H., Andreotti, F., Cooray, N., Chen, O. Y. & De Vos, M. Joint Classification and Prediction CNN Framework for Automatic Sleep Stage Classification. *IEEE Trans. Biomed. Eng.* (2018).

doi:10.1109/TBME.2018.2872652

11. Phan, H., Andreotti, F., Cooray, N., Chen, O. Y. & De Vos, M. SeqSleepNet: End-to-End Hierarchical Recurrent Neural Network for Sequence-to-Sequence Automatic Sleep Staging. *IEEE Trans. Neural Syst. Rehabil. Eng.* (2019). doi:10.1109/TNSRE.2019.2896659
12. Phan, H., Andreotti, F., Cooray, N., Chen, O. Y. & Vos, M. D. Automatic Sleep Stage Classification Using Single-Channel EEG: Learning Sequential Features with Attention-Based Recurrent Neural Networks. *Conf. Proc. IEEE Eng. Med. Biol. Soc.* **2018**, 1452–1455 (2018).
13. Biswal, S. *et al.* Expert-level sleep scoring with deep neural networks. *J. Am. Med. Inform. Assoc.* **25**, 1643–1650 (2018).
14. Sun, H. *et al.* Large-Scale Automated Sleep Staging. *Sleep* **40**, (2017).
